# Obligate movements of an active site-linked surface domain control RNA polymerase elongation and pausing via a Phe-pocket anchor

**DOI:** 10.1101/2021.01.27.428476

**Authors:** Yu Bao, Robert Landick

## Abstract

The catalytic trigger loop (TL) in RNA polymerase (RNAP) alternates between unstructured and helical hairpin conformations to admit and then contact the NTP substrate during transcription. In many bacterial lineages, the TL is interrupted by insertions of 2–5 surface-exposed, sandwich-barrel hybrid motifs (SBHMs) of poorly understood function. The 188-aa, 2-SBHM *E. coli* insertion, called SI3, occupies different locations in halted, NTP-bound, and paused transcription complexes, but its dynamics during active transcription and pausing are undefined. Here we report design, optimization, and use of a Cys-triplet reporter to measure the positional bias of SI3 in different transcription complexes and to determine the effect of restricting SI3 movement on nucleotide addition and pausing. We describe use of H_2_O_2_ as a superior oxidant for RNAP disulfide reporters. NTP binding biases SI3 toward the closed conformation whereas transcriptional pausing biases SI3 toward a swiveled position that inhibits TL folding. We find that SI3 must change location in every round of nucleotide addition and that restricting its movements inhibits both transcript elongation and pausing. These dynamics are modulated by a crucial Phe pocket formed by the junction of the two SBHM domains. This SI3 Phe pocket captures a Phe residue in the RNAP jaw when the TL unfolds, explaining the similar phenotypes of alterations in the jaw and SI3. Our findings establish that SI3 functions by modulating the TL folding to aid transcriptional regulation and to reset secondary channel trafficking in every round of nucleotide addition.

**SIGNIFICANCE:** RNA synthesis by cellular RNA polymerases depends on an active-site component called the trigger loop that oscillates between an unstructured loop that admits NTP substrates and a helical hairpin that positions the NTP in every round of nucleotide addition. In most bacteria, the trigger loop contains a large, surface-exposed insertion module that occupies different positions in halted transcription complexes but whose function during active transcription is unknown. By developing and using a novel disulfide reporter system, we find the insertion module also must alternate between in and out positions for every nucleotide addition, must swivel to a paused position to support regulation, and, in enterobacteria, evolved a “Phe pocket” that captures a key phenylalanine in the out and swivel positions.

## Introduction

Gene expression in all cellular life forms is accomplished by a conserved, multi-subunit RNA polymerase (RNAP) via a highly regulated nucleotide addition cycle (NAC; Fig. 1) that extends RNA transcripts by reaction with DNA-templated nucleoside triphosphates (NTPs). A posttranslocated RNAP first samples NTPs until the correct NTP binds in its active site. A flexible trigger loop (TL) then folds into a helical hairpin called the trigger helices (TH) that is stabilized by contacts to the complementary NTP–DNA pair, causing active-site closure (1-4). The closed active site accelerates by ≥10^4^ S_N_2 displacement of pyrophosphate from the NTP by RNA 3′-hydroxyl-group attack on the *α*-phosphate, resulting in extension of the RNA by one nucleotide (2, 5, 6). Pyrophosphate release, RNA–DNA translocation, and TH unfolding then reset the NAC to the posttranslocated state for the next round of NTP binding. In many bacterial lineages, TL–TH cycling occurs in every round of nucleotide addition despite the presence of a large insertion of 2–5 surface-exposed, sandwich-barrel hybrid motifs (SBHMs) (7-9). The 188-aa, 2-SBHM insertion in *E. coli* RNAP is called sequence insertion 3 (SI3) (10).

**Fig. 1.**
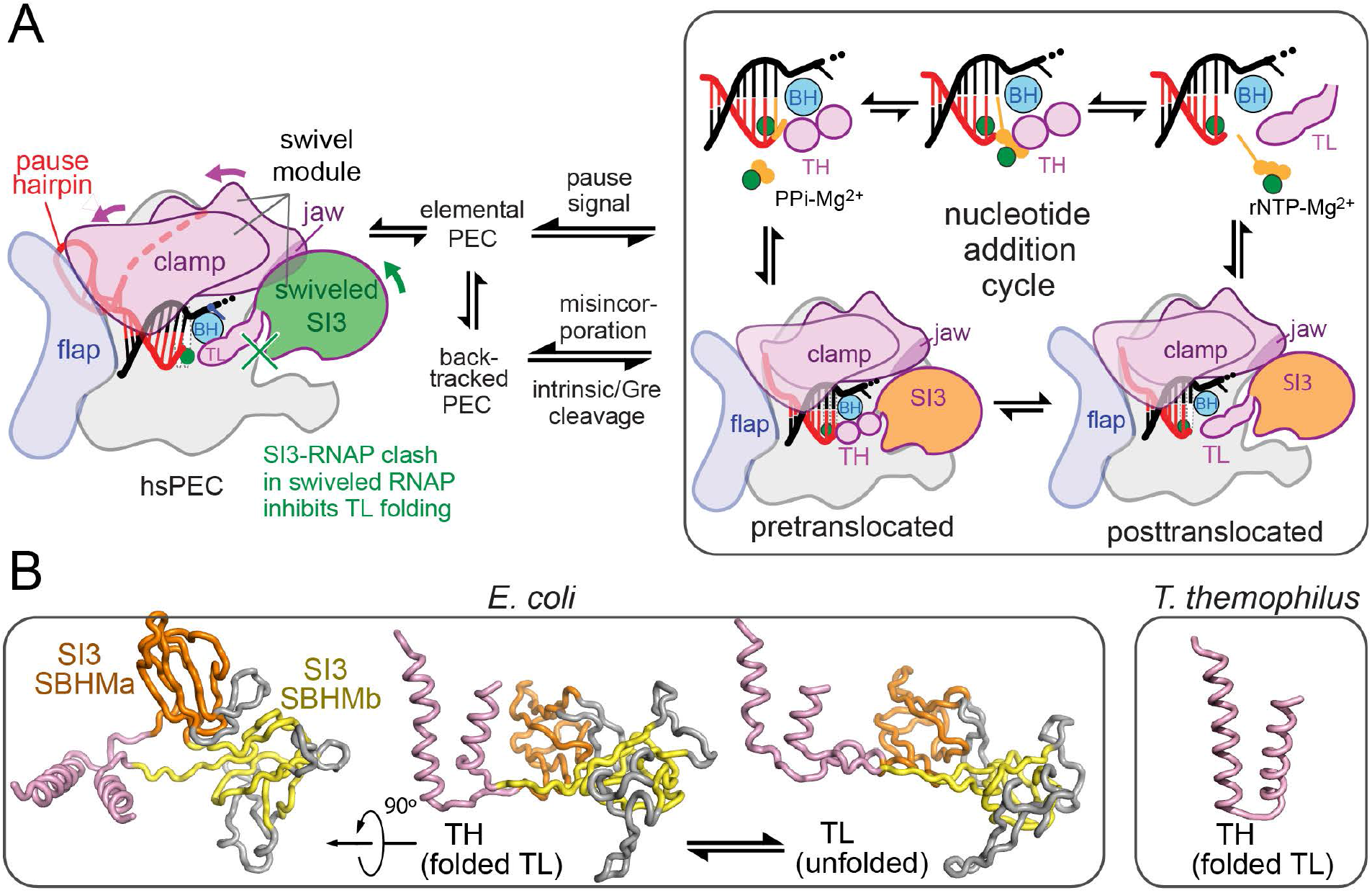
*E. coli* RNAP transcription cycle and SI3. (*A*) The *E. coli* EC alternates between pre- and post-translocated states and between TL and TH states during the nucleotide addition cycle (*right*) but enters alternate conformations in response to pause signals in the DNA and RNA that can slow translocation and induce swiveling or backtracking. SI3 occupies different locations in these states and inhibits TH formation in the swiveled state due to steric clash. (*B*) SI3 in *E. coli* TL (pink) is composed of two SBHM domains (orange and yellow), which include several loops (white) in addition the core SBHM fold. Structures are from pdb 6rh3 (TH) and 6rin (TL) (24). The *T. thermophilus* TH (*right panel*; pdb 2o5j) (2), which lacks a TL sequence insertion, is shown for comparison.

Nucleotide addition is rapid (30–100 nt·s^−1^) at most DNA positions. However, the elongation complex (EC) is controlled in part by pause sequences that halt the EC for ≥1 s by converting it to a paused EC (PEC) (11-13). These transcriptional pauses aid regulation and RNA biogenesis in both prokaryotes and eukaryotes by enabling transcriptional attenuation, transcription–translation coupling, RNA folding, RNA splicing, termination, and antitermination (14-20). Different types of pauses involve distinct PEC conformations, some of which can interconvert (Figure 1A). The elemental pause occurs at specific DNA sequences that inhibit DNA template base loading (i.e., DNA translocation) during the NAC (11). A modest conformational change called ‘swiveling’ and base-pairing energetics may explain slow translocation in the elemental PEC (ePEC), which can occupy multiple states and may rearrange further into backtracked or hairpin-stabilized PECs (21-24). Backtracked PECs are prevalent in eukaryotes, also occur in prokaryotes, and can be triggered by nucleotide misincorporation (25-28). At a hairpin-stabilized pause (e.g., the well-studied *his* pause from the attenuation control region of the *E. coli* histidine biosynthetic operon) (16, 29), a nascent RNA hairpin formed 11– 12 nt from the RNA 3′ end remodels the PEC and stabilizes the swiveled conformation (22, 23). Swiveling involves an ∼4° rotation of a module composed of the clamp, dock, shelf, β′ C-terminal region, jaw, and SI3 (the swivel module). The swivel-module rotation distorts an SI3-complementary depression in the RNAP surface so that RNAP can no longer accommodate the folded TH position of SI3. Backtrack pauses also induce swiveling similar to that seen in *his*PEC (24).

SI3 connects to the apex of the TL–TH via two flexible linkers (7), contacts the β′ jaw in the TL conformation (5), and mediates hairpin-stabilized pausing in *Eco*RNAP by inhibiting TL folding in the swiveled PEC conformation. SI3 and jaw deletions abrogate hairpin-stabilized pausing with no additional effect when combined (5, 30, 31). SI3 shifts to a different location when the TH form, disrupting the SI3–jaw interface (23, 24, 32, 33).

Several aspects of SI3 function remain unclear. Although the TL folds and unfolds in every round of the NAC as expected (34), it is unclear whether SI3 also fluctuates between open and closed locations during rapid transcription or only opens when the EC is halted or paused.

Second, it is unknown if SI3 swiveling is required for pausing or just a consequence of other RNAP conformational changes responsible for pausing. Finally, the structural basis by which the jaw domain modulates SI3 movements is undefined.

To address these questions, we sought ways to exploit engineered disulfide crosslinks both to measure and to test SI3 conformational changes. Previously used methods to form RNAP disulfides proved unable to generate a key SI3 disulfide efficiently, leading us to develop a new approach based on H_2_O_2_ oxidation. The facile formation of stable disulfides with limited damage to RNAP using H_2_O_2_ followed by catalase allowed us to determine SI3 positional biases in different EC states and to analyze how SI3 locations affect transcription and pausing, including in mutant RNAPs that define a role for SI3–jaw interaction.

## RESULTS

### Design and optimization of an SI3 positional reporter – H_2_O_2_ is a superior thiol oxidant

To design disulfide reporters of SI3 location, we examined SI3 residues that changed potential contacts in RNAP structures in which different SI3 positions have been resolved (closed SI3, TH form; open SI3, TL form; swiveled SI3, hairpin-stabilized pause form; Fig. S1*A*) (23, 32, 33). We first sought an SI3 residue that would make different contacts in the closed, open, and swiveled SI3 positions so that potential disulfides created by substitutions could report which location predominates in different EC states. Unfortunately, no SI3 residue exhibits this property. However, the closed and swiveled SI3 states could be distinguished by the location of residues in a β-ribbon-resembling loop in SI3-SBHM1 (β′1046–1061). This loop inserts between the rim helices (RH) and sequence insertion 1 (SI1) such that β′D1051 approaches β′G671 in the closed state, whereas β′D1051 approaches a different part of SI1 (βC267) in the swiveled state. We predicted that Cys substituted for β′D1051 would compete for disulfide formation with Cys residues substituted for β′G671 and βC267 in the closed and swiveled states, respectively (Fig. 2*A,B*).

**Fig. 2.**
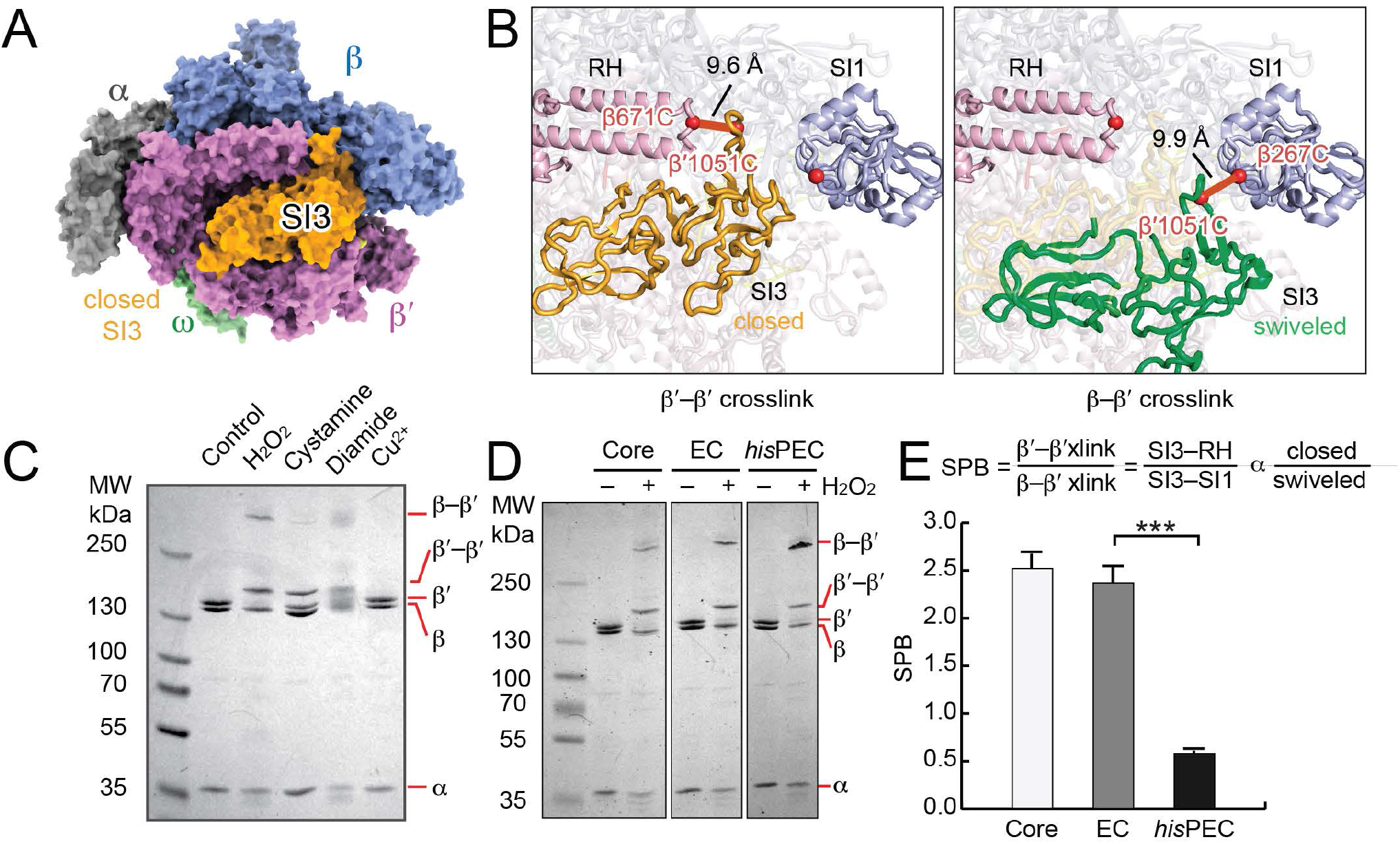
Alternative SI3 locations and detection by a Cys-triplet reporter (CTR). (*A*) RNAP in TH conformation with SI3 in closed position (pdb 4yln) (33). (*B*) C*α*–C*α* distances of residues changed to Cys to generate the CTR (red spheres): β′SI3–β′RH (closed, orange, pdb 4yln) and β′SI3–βSI1 (swiveled, green, pdb 6asx). (32, 33). RH, rim helices; SI1, sequence insertion 1. (*C*) Comparison of ability of different oxidants to form disulfides in the CTR (Materials and Methods). (*D*) Comparison of CTR disulfides formed in apo RNAP core enzyme, an EC, and a paused EC (*his* hsPEC). Both β′–β′and β′–β disulfides are formed in all three complexes. (*E*) SI3 positional bias (SPB) calculated from the ratio of the 2 crosslink species in apo RNAP, the EC and the *his* hsPEC. Results are mean ± SD (n = 3). Unpaired t test, ***p < 0.001.

To test this prediction, we generated an RNAP bearing β′D1051C, β′G671C, and βR267C substitutions. We call this system in which disulfides that report two different conformations compete for formation a Cys-triplet reporter (CTR) (23, 35). Conveniently, the disulfides formed by the SI3 CTR are readily distinguishable because one is internal to the β′ subunit, altering its electrophoretic mobility, and the other links β′ to β, causing a much larger retardation of electrophoretic mobility (Fig. 2*C,D*; Fig. S2).

To date, cystamine (AED), diamide (TMAD), or Cu^2+^ have been used as oxidants to form disulfide bonds in Cys-substituted RNAPs (5, 23, 35-38). However, these oxidants did not work well for the SI3 CTR Fig. 2*C*; Fig. S2*A,B*) and each has disadvantages for probing RNAP conformations. Disulfide formation requires activation of one Cys sulfur by conjugation to the oxidant followed by S_N_2 nucleophilic displacement of the activating group by the second Cys thiolate via a trigonal bipyramidal transition state (Fig. S1*C*). This mechanism can reduce overall crosslink efficiency in two ways (37, 39). First, although high concentrations of oxidant favor disulfide formation electrochemically, some oxidants (e.g., cystamine) also react with the activated intermediate causing accumulation of mixed disulfides in competition with formation of the desired disulfide. Second, steric clash of the conjugated activating group with the surrounding protein environment may inhibit formation of the trigonal bipyramidal geometry in the transition state required for disulfide formation. This latter issue is a particular problem in the complex structure of RNAP where larger or less flexible activating groups often reduce crosslink efficiency. Finally, oxidants can also be disadvantageous because they attack other exposed Cys residues on RNAP and alter its properties (e.g., heavy metals like Cu^2+^ can bind and inactivate enzymes) (40). Such effects may explain why diamide causes smearing of subunit bands during PAGE, complicating quantitation of disulfide bond formation (Fig. 2*C*).

In search of better oxidants for disulfide probes of RNAP conformation, we tested hydrogen peroxide (H_2_O_2_). Compared to cystamine or diamide, H_2_O_2_ generated SI3 disulfides with higher efficiency (>80%) and at relatively high rate (*k*_obs_ = ∼1.5 min^−1^; Fig. S2). H_2_O_2_-induced crosslinks also form in two steps (41). A cysteine thiolate reacts with H_2_O_2_ to form a sulfenic acid (-CSOH) followed by displacement of OH^−^ by a second cysteine thiolate. However, disulfides can also form by conjugation of two sulfenic acids via a thiosulfinate (41), which coupled with minimal steric constraints on the transition state may explain why H_2_O_2_produces disulfide crosslinks with higher efficiency.

The higher efficiency of H_2_O_2_-induced disulfide formation allowed us to create an SI3 CTR in which β′C671 in the RH and βC267 in SI1 compete to form disulfides with β′C1051 in SI3, thus reporting the occupancy ratio of the closed vs. swiveled SI3 conformations (Fig. S1E). In contrast, cystamine was unable to generate the β′C1051-βC267 at useful levels irrespective of RNAP conformation (Fig. 2*C*).

H_2_O_2_ offers an additional advantage over other oxidants. It can be rapidly eliminated from a solution by addition of catalase, which converts H_2_O_2_ to H_2_O and O_2_ with a maximal turnover number of 16,000,000–44,000,000 s^−1^(42). Thus, catalase treatment minimizes oxidative damage to the RNAP once disulfides are formed, allowing *in vitro* transcription or other assays of crosslinked RNAP in the absence of undesired effects of the added oxidant. Catalase treatment also generated clearer bands in SDS-PAGE for crosslink quantitation (compare to diamide, Fig. 2*C*, or H_2_O_2_ without catalase, Fig. S2*F*). Finally, H_2_O_2_ treatment had minimal effects on RNAP activity (Fig. S3). We conclude that H_2_O_2_ is a superior oxidant for disulfide probing of RNAP conformations.

### SI3 samples multiple locations in a resting EC

Using the optimized CTR system, we found that both crosslink species (SI3–RH and SI3–SI1) form in an EC, a paused complex (*his*PEC), and core RNAP (Figs. 2*D*, 2*E*). This result indicates that SI3 samples multiple locations by thermal fluctuation including the closed and swiveled states even in the absence of pause signals or bound NTP substrate. Differences in the ratio of SI3–RH to SI3–SI1 crosslinks should be proportional to changes in the relative occupancies of these states in different transcription complexes (*i*.*e*., proportional to differences in Δ*G*° of the states in different transcription complexes). This view of RNAP as a dynamic system sampling diverse conformational states with different probabilities is consistent with both MD simulations (43) and with the typically lower resolution observed in cryo-EM analyses of RNAP complexes compared to those achievable in globular enzymes (23, 24, 32, 35).

Before using the SI3 CTR to study RNAP, we considered two questions. First, does the formation of both crosslink species by the CTR RNAP reflect non-interconverting conformations of SI3 in separate populations on RNAP? Using RNAPs containing only two Cys substitutions so that only one or the other disulfide could form (i.e., either the SI3–RH or SI3–SI1 disulfide in RNAPs called Cys-pair reporters, CPRs) (37), we found that SI3 conformations are in dynamic equilibrium. Fluctuations in SI3 position allowed each CPR disulfide to form at high efficiency absent conformation with the alternative disulfide (>80%; Fig. S2*E*), which would not be possible if the conformations did not interconvert. Thus, SI3 readily samples the closed or swiveled conformation even in the absence of an NTP substrate or a pause signal.

Second, how is the ratio of SI3–SI1/SI3–RH disulfides in the CTR RNAP related the ratio of the corresponding SI3 conformations? Although we cannot determine this relationship without an independent measure of the SI3 conformations, it seems unlikely that the SI3–RH/SI3–SI1 disulfide ratio and the closed/swiveled conformation ratio are identical. The rate at which each crosslink forms in the CTR RNAP will be a function not only of the ratio of the SI3 conformations but also of the chemical environment around the Cys residues and the probability that the side chains can achieve the transition state sterically. These environmental effects are unlikely to be identical for the competing disulfides. Nonetheless, because the SI3 states are dynamically interconverting, a change in the SI3–RH/SI3–SI1 disulfide ratio should be proportional to changes in the bias of the SI3 positional equilibrium for a given RNAP or EC state. We defined ‘SI3 Positional Bias’ (SPB) as the ratio of the SI3–RH/SI3–SI1 disulfides and a relative measure of closed/swiveled SI3 occupancy (Fig. 2*E*). A higher SPB indicates that SI3 is more biased toward the closed position and a lower SPB indicates SI3 is more biased toward the swiveled position.

### Cognate NTP binding stabilizes SI3 shift to closed conformation

We first sought to explore how SI3 position changes upon substrate binding. Structural studies suggest that SI3 favors the closed position when cognate NTP is bound by an EC (24, 33), conditions known to stabilize TH formation to promote catalysis (23, 32, 33, 44). However, the positional bias of SI3 upon binding of cognate or noncognate NTPs in physiologic conditions where SI3 is dynamic remains untested. To ask how NTP binding affects SI3 location, we used SI3-CTR RNAP and an RNA–DNA scaffold to form an EC that allowed NTP binding but not nucleotide addition. This scaffold contained complementary 50mer DNA strands and a 19-nt RNA that was reacted with ATP to form A20 ECs or with 3′dATP to form dA20 ECs (Fig. 3*A*; Fig. S4*A*). Upon oxidation, A20 EC gave an SPB of 1.8 ± 0.1, similar to U19 (1.2 ± 0.1), but the SPB shifted to 5.9 ± 0.5 upon addition of the unreactive CTP analog CMPCPP (Fig. 3*B*). This shift in SBP indicates that SI3 increases occupancy of the closed position by a factor of ∼3.3 upon cognate NTP binding. Similarly, when CTP was added to dA20 EC, which lacked the 3′ OH required for S_N_2 attack during nucleotide incorporation, SPB shifted from 2.4 ± 0.2 to 6.7 ± 0.9, an increase of ∼2.8x. The slightly higher SPB of dA20 EC vs. A20 EC is consistent with prior observations that a 3′ deoxy substitution favors the pretranslocated register, which favors TL folding (45).

**Fig. 3.**
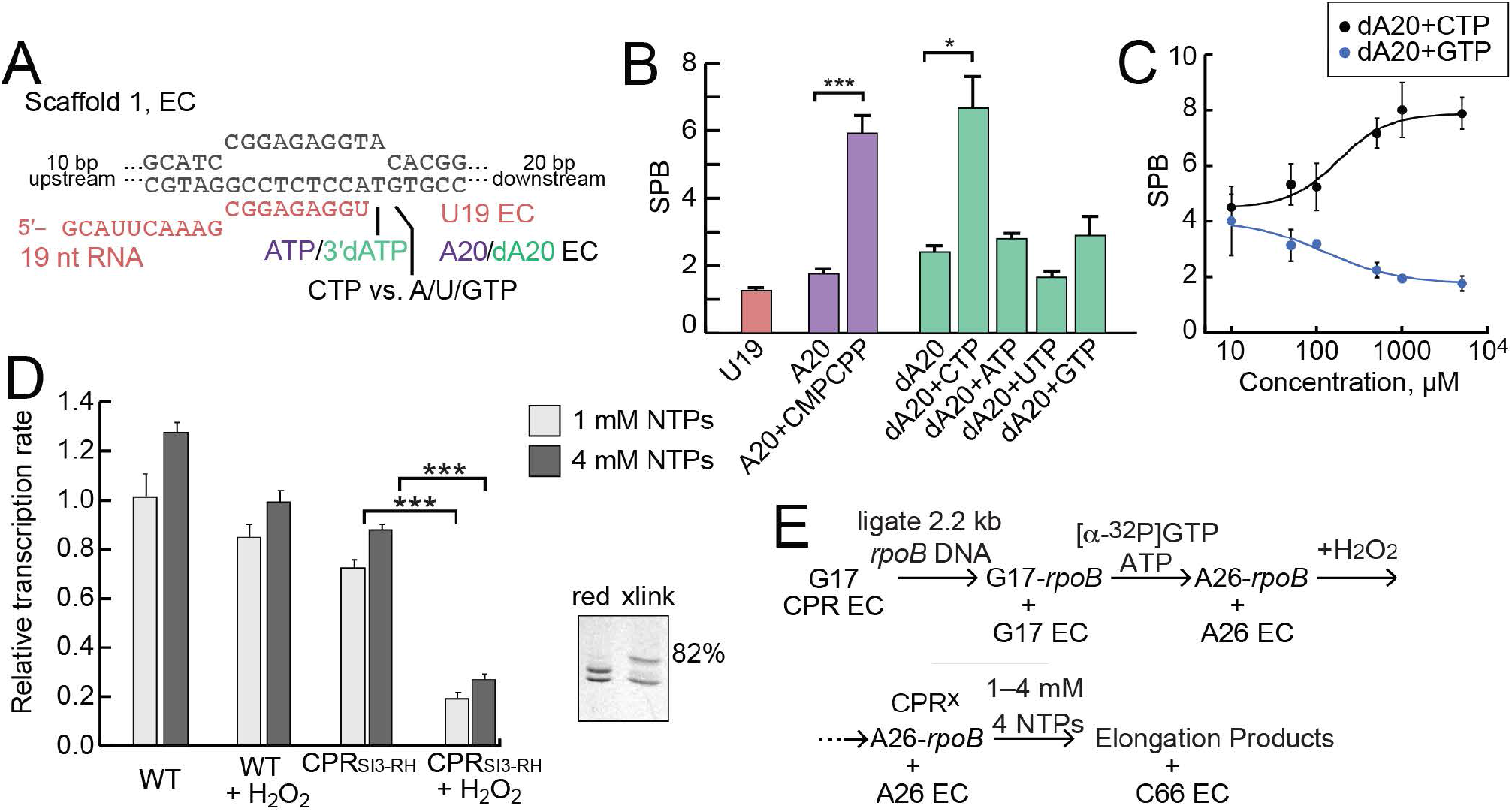
SI3 is biased toward the closed position by cognate but not noncognate NTPs and inhibits transcript elongation when crosslinked in the closed position. (*A*) RNA–DNA scaffold used to reconstitute ECs. Either ATP or 3′ deoxy ATP (3′dATP) can be incorporated at position 20 (see also Fig. S4). 3′dA20 EC supports NTP binding but not nucleotide addition. (*B*) SPB of A20 EC, 3′dA20 EC, and ECs incubated with different NTPs (at 100 µM each NTP). (*C*) SPB of 3′dA20 EC incubated with different concentrations of cognate CTP or non-cognate GTP. (*D*) Average elongation rates (relative to WT RNAP in reducing condition) at 1 mM each NTP or 4 mM each NTP. Data represent mean ± SD (n = 3). Unpaired t test, ***p < 0.001. Inset, a typical assay of crosslink efficiency (82% of CPR_SI3-RH_ was crosslinked by 5 mM H_2_O_2_). (*E*) Schematic of ligated-scaffold transcription assay. The ligation reaction is incomplete, but only products longer than C66 are used to estimate elongation rate (see Fig. S5 and *Materials and Methods*).

We next asked how cognate vs. noncognate NTP would affect SI3 positional bias. At 100 µM, noncognate ATP and GTP marginally increased dA20 SBP whereas UTP decreased SPB possibly because rU-dG base-pairing disfavors TL folding (Fig. 3*B*). Similarly, at higher concentration (5 mM) noncognate GTP decreased SPB to 1.8 ± 0.3 with a concentration dependence similar to that by which cognate CTP raised SPB from 4.5 ± 0.5 at 10 µM to 7.9 ± 0.6 at 5 mM (Fig. 3*C*). These concentration profiles with half-maximal but opposite effects at ∼180 µM are similar to previous measurements of NTP binding to 3′deoxy ECs (37, 45) and are consistent with a pre-insertion model for NTP selection in which NTP binding is indiscriminate and substrate discrimination arises principally from the ability of a complementary rNTP-dNMP base pairing to support TL folding (3, 46). The decrease in SPB caused by GTP suggests that either non-Watson–Crick base pairing or NTP binding in the E site may decrease occupancy of the TH state due of steric clash with TL side chains. Taken together, these results suggest that cognate NTPs shift SI3 positional bias toward the closed position, likely in concert with TL folding, whereas noncognate NTPs may shift SI3 away from the closed positions, likely linked to decreased TH formation. Thus, the energetics of SI3 positioning may aid in NTP discrimination by modulating TL folding, which directly affects NTP discrimination (47-49), although effects of SI3 on NTP selectivity remain to be investigated in detail.

### Nucleotide addition cycling requires SI3 open–close cycling

A key question is whether SI3 moves back and forth between the closed and open position in every round of the NAC. Although the TL clearly undergoes folding unfolding cycles in each round of nucleotide addition (34) and blocking TH unfolding by disulfide crosslinking inhibits rapid transcription (37), it is unknown if SI3 also moves in every round of the NAC or might only shift to the open or swiveled state when an EC halts or pauses. To determine whether SI3 movement is required for rapid nucleotide addition, we asked how biasing SI3 to the closed position with a disulfide would affect transcription rate during multi-round nucleotide addition. For this purpose, we used a previously described ligated-scaffold transcription assay that enables transcription by reconstituted ECs over multi-kb of DNA (50). We ligated a 2.2 kb DNA from the *E. coli rpoB* gene to reconstituted CPR_SI3–RH_ ECs and then measured transcript elongation at 1 mM each of all four NTPs with and without the SI3–RH disulfide (Fig. 3 *D* and *E*). Biasing SI3 to the closed position with the disulfide allowed transcript elongation but at an average rate that was slower by a factor of ≥3.7 compared to the non-crosslinked EC (Fig. 3*D*; Fig. S5). Pausing by the SI3-crosslinked RNAP was enhanced at some locations (e.g., at G31, 5 nt downstream of A26) and only a few ECs were able to extend past the point of ligation (+66; Fig. S5*B*). The inhibition of multi-round transcript elongation when SI3 is crosslinked in the closed position is unlikely due only to reduced affinity of ECs for substrate NTP, since increasing Mg^2+^·NTP concentrations 4-fold to 4 mM each had little effect on transcription rate (Figure 3*D*).

These results indicate that SI3 must ordinarily alternate between the closed and open positions in each round of nucleotide addition rather than remaining closed while accommodating TL–TH cycling (note that biasing SI3 open is even more inhibitory as expected based on a TH-containing EC structure; see below and Fig. 5). Biasing SI3 to the closed position likely influences transcription in multiple ways. First, it should favor TH formation, which narrows the RNAP secondary channel and should restrict NTP access to active site. Second, the closed SI3– TH may stabilize the pretranslocated state during the NAC, which could slow transcription by inhibiting translocation as observed for a TH-stabilizing crosslink (37). Third, the closed SI3–TH appeared to strengthen sequence-specific pausing at certain sites (e.g., the pause sites evident in Fig. S5*B*). The SI3–RH crosslink must allow some extent of TH–TL interconversion, however, since it decreased transcription rate by less than a factor of ten whereas deletion of the TL inhibits transcription rate by ≥10^4^ and a direct stabilization of the TH inhibits transcription rate by factors of ≥10–200 (2, 5, 6, 44). We conclude that alternation of SI3 between open and closed locations likely occurs in every round of nucleotide addition, and that inhibition of these oscillations explains the effect of the SI3–RH disulfide.

**Fig. 5.**
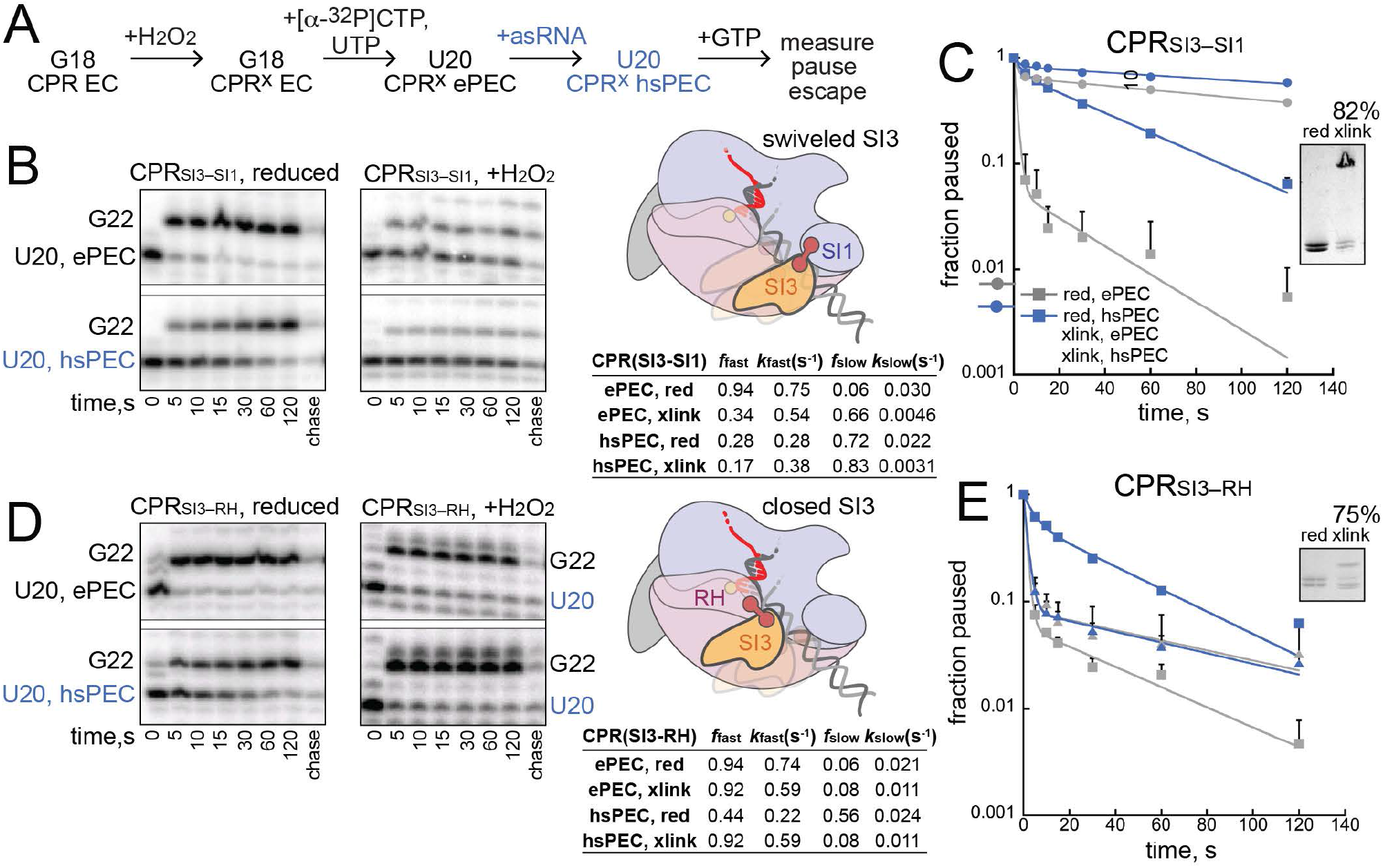
Biasing SI3 to swiveled or closed positions with disulfides establishes that SI3 swiveling is sufficient and necessary for hairpin-stabilized pausing. (*A*) Reaction schematic to test effects of SI3 disulfides on pausing. (*B*),(*D*) RNAs formed during pause assays, schematics of SI3 disulfides tested, and pause kinetics. *f*_fast_ and *f*_slow_ are the fractions of pauses in faster and slower states when fit to a two-exponential equation and *k*_fast_ and *k*_slow_ are the corresponding rate constants (*Materials and Methods*). A complete set of kinetic fitting parameters with errors is in Table S2. (*C*),(*E*) Fraction pause RNAs as a function of time under different conditions.

### SI3 location bias shifts in paused ECs

We next examined the effect of RNAP pausing on the location bias of SI3 using the well-characterized paused ECs (PECs) that form at U102 in the attenuation-controlled leader region of the *E. coli his* biosynthetic leader region (*his*PECs) (22, 23, 29). RNAP forms either ePECs in the absence of a nascent RNA pause hairpin or hairpin-stabilized PECs (hsPECs) when a nascent RNA structure forms in the RNA exit channel 11 nt from the RNA 3′ end. We examined ECs (G18), ePECs (U20), and hsPECs (C19 and U20) reconstituted with CTR RNAP on the same scaffold using an antisense RNA to form the pause hairpin, limited nucleotide addition to advance G18 ECs to C19 (with CTP) and U20 (with CTP and UTP), and either a complementary or noncomplementary (bubble) NT DNA strand (Fig. 4*A*). We observed higher SPBs on the bubble-containing ECs and PECs, but the same overall trends in SI3 positional bias (Figs. 4 *B,C*). The higher SPBs on bubble-containing complexes may reflect lesser thermal fluctuations of SI3 when translocation is constrained by the noncomplementary bubble, allowing greater formation of the closed SI3 crosslink. In both types of complexes, however, the ePEC exhibited a modest shift toward the swiveled conformation relative to G18 and C19 ECs whereas the hsPEC increased the bias toward the swiveled conformation more dramatically (Fig. 4 *B,C*).

**Fig. 4.**
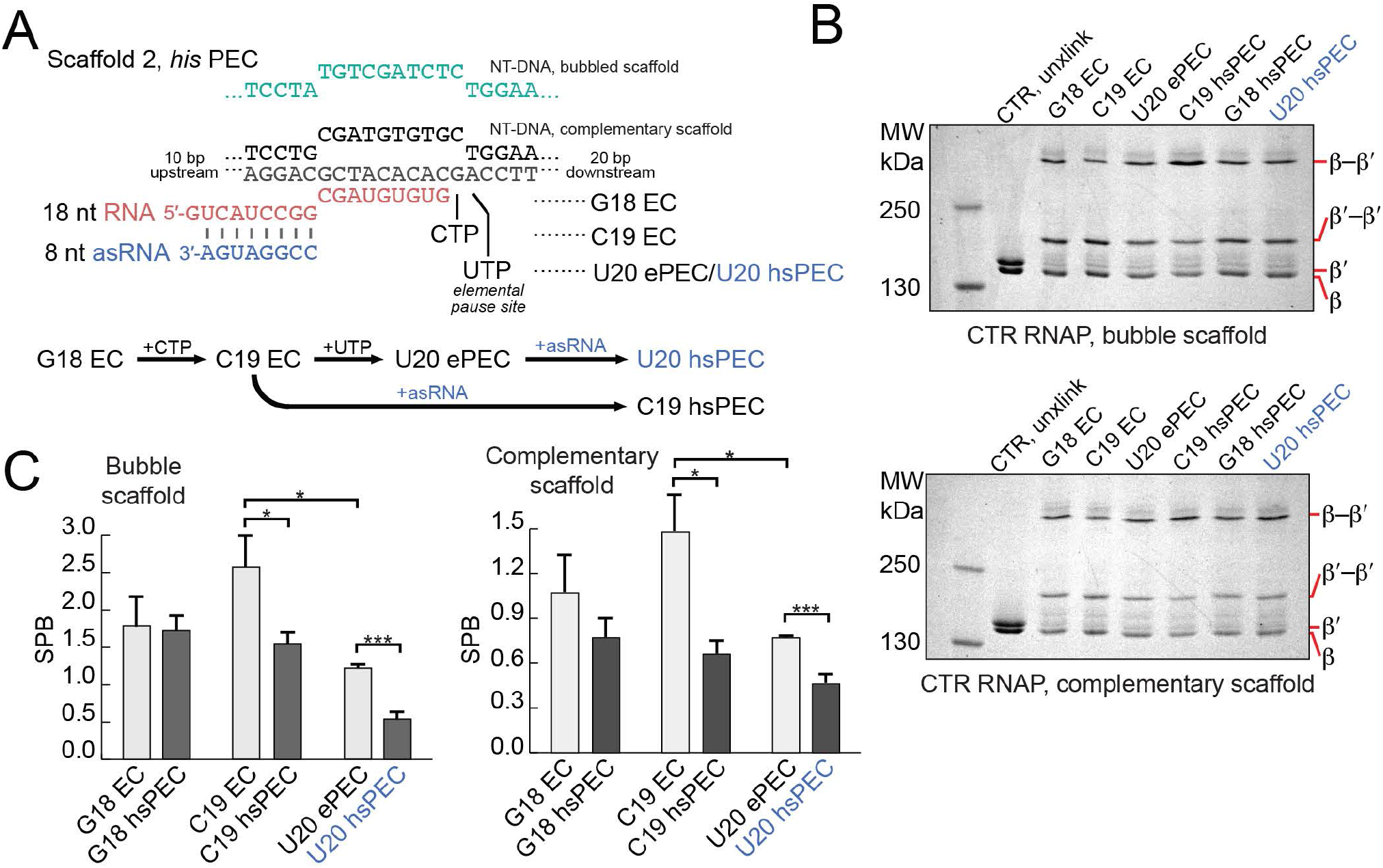
SI3 becomes biased to the swiveled position as RNAP in the *his* ePEC and hsPEC. (*A*) RNA–DNA scaffold used to reconstitute *his* PECs. G18 EC reacts with CTP and UTP to form C19 EC and U20 ePEC, respectively (See also Figure S3). Addition of an asRNA mimics formation of the *his* pause hairpin and forms the *his* hsPEC. (*B*) SDS-PAGE images of CTR crosslinks formed on complementary and bubble scaffolds of his ECs and PECs. (*C*) SPB of *his* ECs and PECs. Data represent mean ± SD (n = 3). Unpaired t test, ***p < 0.001, *p < 0.05.

These changes in SI3 positional bias for the *his* elemental and hairpin-stabilized pauses confirm that SI3 swiveling imaged by cryoEM of PECs reflects changes in the dynamics of SI3 in physiological condition and that the extent of swiveling correlates with pause duration since SI3 in the *his* ePEC is less biased toward the swiveled state than in longer-lived *his* hsPEC. Hairpin promotion of swiveling is evident at C19 in addition to U20, consistent with prior observations of that the *his* pause hairpin can form in the pause –1 complex (51, 52); this result also supports the conclusion exit-channel RNA duplexes directly promote SI3 swiveling. A shift in SI3 toward the closed conformation in C19 ECs relative to G18 ECs also was evident (Fig. 4 *B,C*), consistent with prior observations that 3′ C favors the pretranslocated state, 3′ G favors the posttranslocated state, and a pretranslocated EC stabilizes the TH vs. TL conformation (45, 53).

### SI3 swiveling contributes to the hairpin-stabilized pause

Although both cryoEM and our CTR data support a causal role for SI3 swiveling in pausing, a direct experimental test by preventing or promoting swiveling is needed to establish causality. To perform this test, we next used CPR_SI3–SI1_ and CPR_SI3–RH_ RNAPs to probe the effects on *his* pausing of strongly biasing SI3 to the swiveled or closed conformation, respectively. The SI3– SI1, swiveled crosslink was formed at G18 with high efficiency (∼82%) prior to C19 radio-labeling and U20 extension (Fig. 5*A*). Addition of GTP to the crosslinked PECs enabled measurement of the pause escape rate. Without crosslinking, the majority of the ePEC (94 ± 2%) rapidly escaped the weak elemental pause (*k*_ePEC, red_ = 0.75 s^−1^) whereas the majority of the hsPEC (72 ± 2%) generated after asRNA addition escaped the hairpin-stabilized pause a rate 34 times slower (*k*_hsPEC, red_ = 0.022 s^−1^; Fig. 5 *B,C*). When CPR_SI3–SI1_ RNAP was crosslinked to favor the swiveled SI3 conformation, the elemental pause escape rate was significantly slowed by a factor of >170 (*k*_ePECx_=0.0046 s^−1^), even slower than that of the uncrosslinked hsPEC. Addition of asRNA to the SI3–SI1-crosslinked U20 EC further decreased the pause escape rate by a factor of only ∼1.5 compared to the 34-fold effect the uncrosslinked EC (*k*_hsPECx_ = 0.0031 s^−1^). These results indicate that artificially biasing SI3 toward the swiveled position greatly increases pause dwell time even in the absence of an exit channel RNA duplex. Nearly all the effect of the pause hairpin was recapitulated by the swivel-biasing crosslink, suggesting that practically all the effect of hairpin is mediated by stabilizing the swiveled SI3 conformation. We conclude that biasing SI3 to the swiveled position strongly inhibits TL folding, thus obviating the swivel-inducing effect the exit-channel RNA duplex created by asRNA addition.

To test the opposite prediction that inhibiting SI3 swiveling with the SI3–RH crosslink should also obviate the effect of an exit-channel RNA duplex, we measured pause kinetics for the *his* PEC formed with CPR_SI3–RH_ RNAP with and without addition of the asRNA. As predicted for the hsPEC, formation of the SI3–RH crosslink to favor the SI3 closed position greatly reduced pausing and essentially eliminated the effect of the exit channel duplex (compare pausing profiles for the crosslinked ePEC and hsPEC, Fig. 5 *D,E*). Biasing SI3 toward the closed conformation either prevented the exit-channel duplex from inducing swiveling or prevented the duplex from forming (e.g., if swiveling is required to accommodate the duplex in the exit channel). These results support the view that induction of SI3 swiveling and consequent inhibition of TL folding is the mechanism by which exit-channel RNA duplexes increase pause dwell time.

In contrast for the *his* ePEC, formation of the SI3–RH crosslink to favor the SI3 closed position modestly increased pausing, contrary to the expectation that swiveling aids elemental pausing (Fig. 5*D,E*). However, this increase in pausing is consistent with stimulation of pausing at some sites observed when crosslinked CPR_SI3–RH_ RNAP transcribed over a long DNA template (Fig. S5*B*). We conjecture that stabilizing SI3 in the closed conformation may stabilized the pretranslocated register of the ePEC and could increase pausing at sites where the translocation is naturally slow.

To summarize, the pausing behavior of PECs with SI3 biased toward either the closed or swiveled position strongly supports the view that swiveling stabilized by an exit-channel duplex or simply in response to RNA and DNA sequences is a direct cause of transcriptional pausing, likely by inhibiting the transition from the half-translocated (RNA-only) to fully translocated state in the NAC. Stabilization of the TH by biasing SI3 toward the closed state also appears capable of stimulating pausing.

### An SI3 Phe pocket captures the RNAP jaw to tune TL folding, pausing, and catalysis

Previous biochemical studies suggest that the β′ jaw domain participates in pausing via interdependent effects with SI3 (5, 30, 31), but the basis of jaw–SI3 interaction has not been defined. We examined the SI3–jaw interface in cryoEM structures of ECs and the *his*PEC (23, 24, 32) and observed a hydrophobic alcove on the SI3 surface (a Phe pocket) that appeared to accommodate the Phe side chain of a jaw residue (β′F1199; Fig. 6*A*). However, when SI3 is in the closed position, this SI3–jaw interaction is disrupted and F1199 no longer resides in the pocket (e.g., when NTP is bound by an EC; Fig. 6*B*) (24). To test whether the jaw F1199–SI3 Phe pocket interaction inhibits movement of SI3 to the closed position, we generated RNAPs bearing a β′F1199A substitution and used them for CTR and pausing assays. We found that SI3 became more biased toward the closed position in both ECs and hsPECs when F1199 was changed to Ala, indicating that the F1199–Phe pocket interaction stabilizes jaw–SI3 interaction and that its disruption decreases the thermodynamic stabilities of SI3 locations in which the interaction occurs (i.e., open and swiveled SI3; Fig. 6*C*). Pause assays using β′F1199A RNAP revealed a reduction in hairpin-stabilized pausing, consistent with the model that SI3 swiveling, stabilized by F1199–Phe pocket interaction, is required for hairpin-stabilized pause (Figure 6*D*). These results suggest that the jaw provides a docking site for SI3 via a Phe pocket interaction that modulates SI3 movements during both the NAC and transcriptional pausing.

**Fig. 6.**
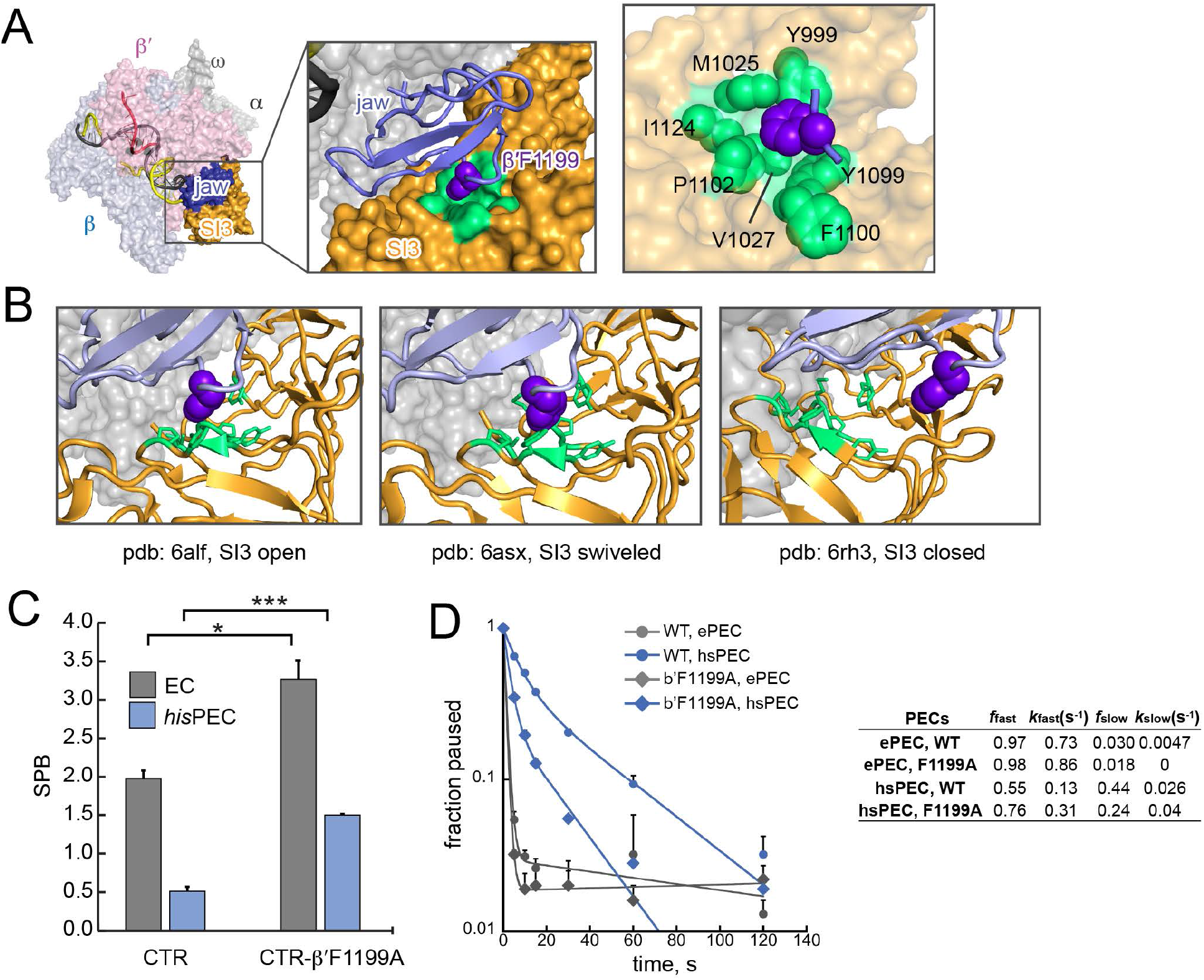
Capture of β′F1199 in the RNAP jaw by a Phe pocket in SI3 modulates SI3 movement. (*A*) Phe pocket interaction in SI3 open position (pdb 6alf) (32). Light green, the hydrophobic pocket in SI3. Purple, β′F1199. (*B*) Phe pocket interaction and disruption in open (pdb: 6alf), swiveled (6asx), and closed (6rh3) ECs or PEC (23, 24, 32). (*C*) CTR assay of wild-type and β′F1199A RNAPs bound to EC and *his*PEC scaffolds. Unpaired t test, ***p < 0.001, *p < 0.05. (*D*) Pause RNA as a function of time after mixing with GTP in reducing conditions and derived pausing kinetic parameters for CTR RNAP and CTR-β′F1199A RNAP. β′F1199A decreased hairpin-stabilized pausing by a factor of ∼2.5. See Fig. 5 legend for kinetic parameter definitions.

## DISCUSSION

We report new insights into the function of SI3, an unusual modulator of RNAP catalytic function, by exploiting sulfhydryls placed at *E. coli* RNAP locations that alternate proximity when SI3 changes location and H_2_O_2_–catalase as a superior disulfide-bond-formation system to actuate disulfide reporters of RNAP conformation: (*i*) SI3 fluctuates among a range of locations from closed to swiveled in most ECs with a predominate location determined by interactions in the EC; (*ii*) SI3 ordinarily cycles between preferentially closed and open positions in every round of nucleotide addition accompanying TL folding–unfolding; (*iii*) SI3 swiveling is a direct cause of prolonged transcriptional pausing; and (*iv*) the open and swiveled conformations of SI3 are stabilized by capture of a Phe residue in the RNAP jaw domain by a Phe-pocket formed at the interface between the two SBHM domains in SI3. These findings have important implications for the function and evolution of SI3-like domains in bacterial RNAPs.

### SI3 positional cycling resets RNAP for timely regulation during transcript elongation

Structural analyses capture SI3 in the open position or as disordered except when active-site bound NTP stabilizes SI3 in a closed position (24, 32, 33, 54) or pausing favors a swiveled position (22-24), but it has been unclear if SI3 actually cycles between closed and open locations in concert with TL folding and unfolding during rapid nucleotide addition. The location of the open SI3 on the jaw domain stabilized by the Phe-pocket interaction appears structurally incompatible with TH formation because the linkers connecting SI3 to the TL are too short to accommodate the TH position without SI3 movement. The opposite is not necessarily true, however, raising the possibility that SI3 could remain closed during rapid TL–TH cycling and nucleotide addition and possibly open only at pause sites or in response to other regulatory events. Our findings contradict this view, since stabilizing the closed SI3 strongly inhibits rapid nucleotide addition (Fig. 3*D*). Instead, they support a view in which SI3 fluctuates among many possible locations including closed, open, and swiveled in most ECs, with strong biases favoring the closed state when substrate binding stabilizes TH formation, the open state after PPi release and translocation, and the swiveled state in paused ECs (Fig. 4*D*; Fig. 7).

**Fig. 7.**
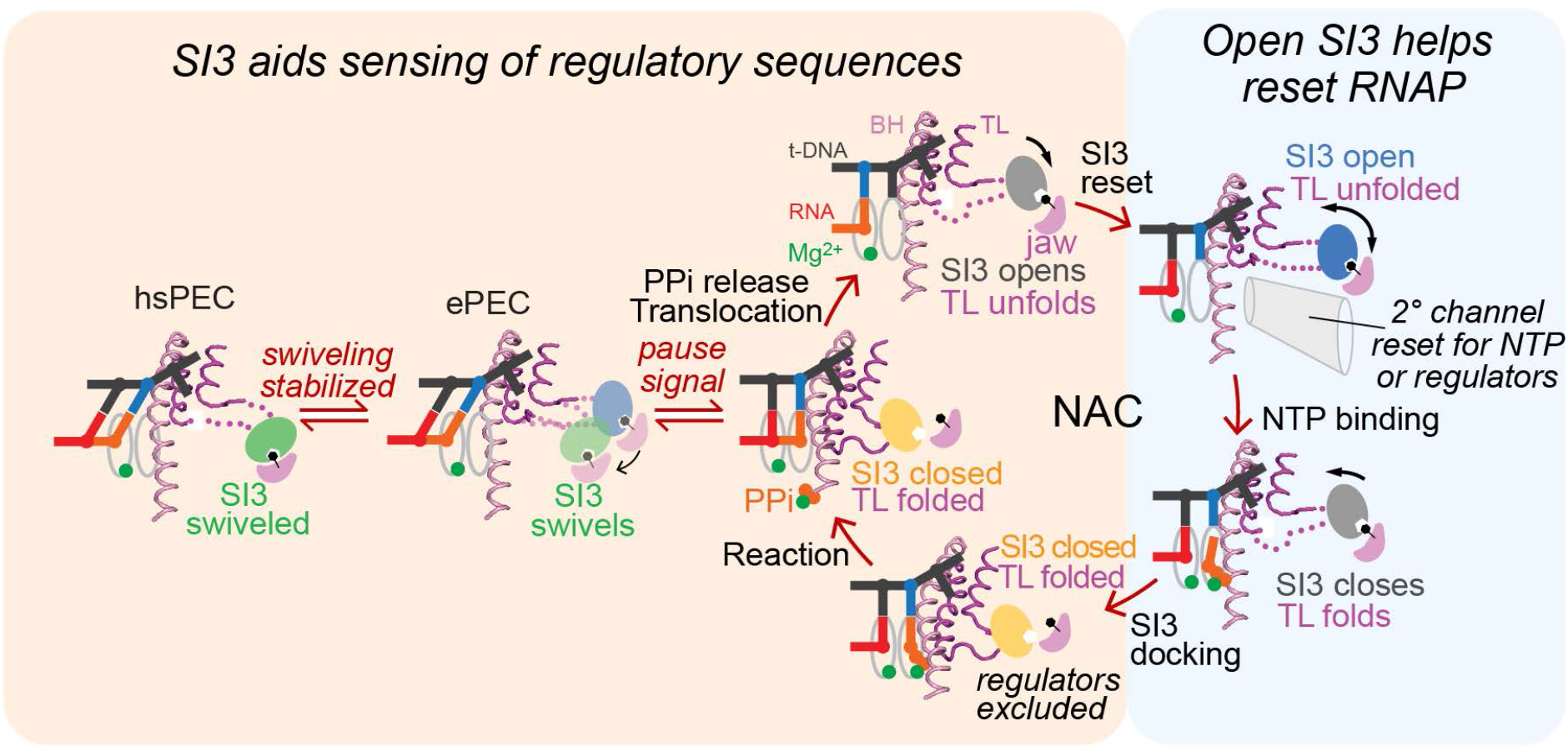
SI3 states during RNA chain elongation and pausing aid transcription. In an active posttranslocated EC with an unfolded TL (NAC, top left), SI3 docks on the jaw domain but thermal fluctuation enables SI3 to sample among different positions. In this state, the secondary channel (2° channel) is open and can admit NTPs or regulators. Binding of cognate NTP then stabilizes the folded TH with SI3 shifted in the closed position. In this location, SI3 will clear the secondary channel of regulators and, together with the TH, close the active site, which promotes the nucleotide addition reaction. After nucleotide addition, the EC either undergoes translocation with PPi release, TL unfolding, and SI3 movement back to the open position or undergoes a transition to the elemental paused state. In response to RNA–DNA sequences, the ePEC can either isomerize back to the elongation pathway (back to the NAC) or convert to either a backtracked PEC or a hairpin-stabilized PEC in which swiveling becomes stabilizes for the pause hairpin.

This picture of a fluctuating SI3 domain that cycles between predominately closed or open positions in concert with the nucleotide addition cycle suggests that SI3 could play an important regulatory function by resetting the RNAP secondary channel and possibly the overall the state of the EC in every round of nucleotide addition. Resetting the secondary channel may be important because multiple secondary channel-binding regulators compete to occupy the secondary channel when SI3 is in the open position, including GreA, GreB, TraR, Rnk, and DksA in *E. coli* and additional regulators in other bacteria (55). When SI3 shifts to the closed position, steric clash would preclude secondary-channel binding by these regulators. NTPs also must enter the active site through the secondary channel but appear unable to pass through when the channel is occupied by a secondary channel-binding factor. By cycling between open and closed locations in every round of nucleotide addition, SI3 effectively resets the secondary channel to a naïve state when it moves from the closed position that excludes the regulators to the open position that allows both NTPs and regulators to compete based only new state of the EC after the last round of nucleotide addition. This resetting function of SI3 could aid efficient regulatory responses of the EC based on its current state.

SI3 cycling also could contribute to resetting the entire EC in every round of nucleotide addition. A long-standing discussion has considered whether RNAP can occupy persistent conformational states over multiple rounds of nucleotide addition; such states might, for instance, be resistant to termination or possess altered catalytic properties independent of bound regulators (56, 57). Although not definitive, cycling of SI3 in every round of nucleotide addition appears inconsistent with the existence of persistent RNAP conformational states since SI3 resets to a state able to transiently occupy multiple locations in every round of nucleotide addition.

### SI3 augments the regulatory capacity of RNAP by enhancing transcriptional pausing

Our results establish that biasing SI3 location toward swiveling with a disulfide bond directly enhances transcriptional pausing, presumably by inhibiting TL folding, stabilizing the half-translocated state observed in the *his* ePEC and hairpin-stabilized PEC, or both (Figs. 1 and 5). Intriguingly, we also observed that some pauses may be increased when SI3 is biased toward the closed conformation by a disulfide bond (Fig. S5*B*). Possibly, this increased pausing by the closed CPR RNAP occurs at sites where PECs can be stabilized in the pretranslocated state. Strikingly, however, the closed CPR RNAP was unaffected by pause hairpin formation at the *his* pause site, both confirming the role of swiveling in pause hairpin action and showing that SI3 affects different classes of pause mechanisms in different ways.

These results suggest that an important function of SI3 could be to increase the potential for sequence-dependent regulation of transcript elongation by sensitizing RNAP to different modes of pausing. The presence in RNAP of a TL insertion able to assume different conformational states increases the conformational complexity of RNAP. By way of its insertion directly into the TL, the conformation of SI3 can affect catalysis. By virtue of being surface exposed, it is possible, although as yet undemonstrated, that solution conditions could affect SI3 conformational states. Thus, by fluctuating among various positions, SI3 may increase the ways that DNA and RNA sequence can bias RNAP to meta-stable pause sites stabilized in part by SI3 contacts or clashes in particular RNAP conformations. In this way, SI3 increases the regulatory capacity of RNAP to respond to transcriptional pausing signals.

### SI3-like insertions are widespread but the Phe-pocket is specific to γ-proteobacteria

Both resetting secondary-channel trafficking and increasing regulatory capacity may explain in part how SI3, and TL insertions more generally, were acquired during evolution of bacterial RNAPs. However, several interesting questions about SI3 evolution remain unanswered. All known TL insertions consist of two or more repeats of SBHM domains (8, 9), but their distribution among bacterial lineages begs the question of whether some insertions in different lineages arose independently or if all TL insertions are evolutionarily related (Fig. S6*A*). TL insertions are present in most bacterial lineages but the lineages in which they are absent vs. present do not cluster together (58), raising the possibility these insertions arose independently. Most notably, insertions (of five SBHM domains) are present in cyanobacteria even though they are absent in most evolutionarily nearby lineages. The SBHM fold is present elsewhere in the RNAP large subunits, including as a universally conserved feature called the flap or wall in bacterial and eukaryotic RNAPs, respectively (8). It is possible that unusual genetic rearrangements that arise over time in large, rapidly growing bacterial populations may have led to duplications of the SBHM fold and independent selection of these TL insertions in RNAP in different lineages.

Regardless of evolutionary history, another question is whether TL insertions are related to another feature of transcriptional regulation that differs in diverse bacteria. Although our understanding of the diversity of regulatory mechanisms in diverse bacteria remains primitive and in great need of expanded study, two distinct themes in bacterial regulation of transcript elongation have recently emerged. First, the universal elongation factor NusG displays markedly different effects in different lineages, stimulating pausing in some like firmicutes and actinobacteria (based *B. subtilis* and *M. tuberculosis*) whereas it suppresses pausing in proteobacteria (based on *E. coli*) (59-61). Second, transcription and translation are coupled in some bacterial lineages like *E. coli* but not coupled in others like *B. subtilis*, with related large differences in the role of Rho-dependent termination in mRNA quality control (62). However, the distribution of TL insertions among bacterial lineages appears to be unrelated to the patterns of either NusG pause effect or transcription–translation coupling among different bacterial lineages (59, 62). Thus, selection for TL insertions appears to have operated independently of these other major themes in the bacterial regulation of transcript elongation.

One possibility is that a major selective pressure for generation or retention of TL insertions is simply to increase the evolvability of RNAP to adapt to diverse environments and stresses (63). In other words, the presence of a TL insertion increases the ways in which the regulatory responses of RNAP can be altered by mutation in response to different selective pressures. Even without mutation, a TL insertion may RNAP easier to regulate by increasing the available conformations that impact catalysis. Especially in bacteria whose range of environmental niches and adaptive responses is huge, an ability to change regulatory responses simply by changing the properties and interactions of a TL insertion may be significant.

One line of evidence supporting selection for evolvability of TL insertions comes from our discovery of the role of the Phe pocket in modulating the function of SI3 in *E. coli* (Fig. 6). Although the sequences that comprise the Phe pocket and the jaw Phe residue itself are well conserved among *γ*-proteobacteria, these residues are not conserved in other bacterial lineages in which TL insertions are present (Figure S6*B*). Indeed, the jaw Phe residue is more commonly His in other TL insertion-bearing lineages. This observation suggests the crucial Phe pocket evolved to modulate SI3 function in *γ*-proteobacteria, whereas other interactions of TL insertions, as yet uncharacterized, may predominate in other lineages. Thus, our current findings highlight two important questions for future study: (*i*) what specific regulatory features are conferred on SI3 by the Phe-pocket interaction; and (*ii*) to what extent do locations and interactions similar to those found for SI3 operate in other TL insertions versus lineage-specific evolution of TL-insertion locations and interactions?

## MATERIALS AND METHODS

Further details are provided in the Supplemental Materials and Methods.

### Reagents and materials

Plasmids and oligonucleotides used in this study are listed in Table S1. The CTR, CPR, and RNAPs were purified as described previously (5, 12, 21) using expression plasmids derived from pRM756 with a His_10_-ppx tag at the C terminus of β′ or, for variants with substitutions in β, from pRM843 with a His_10_-ppx tag at the N terminus of β and a Strep tag at the C terminus of β′ subunit.

### Transcription crosslinking assays

ECs and PECs were reconstituted and assayed for pausing an elongation as described previously (21). To induce disulfide bond crosslinking, H_2_O_2_ was added to 10 µL of CTR or CPR RNAPs or ECs at final concentrations of 5 mM H_2_O_2_ and 2–2.5 µM RNAP or EC and the mixtures were incubated for 15 min at 37 °C. To remove excess H_2_O_2_, 1 µL of 2 U/µL catalase (bovine, Sigma-Aldrich) was then added and incubation was continued for 5 min at room temperature. The ligation–elongation assay using a 2.2 kb DNA (*E. coli rpoB*[216–2459]) with StyI-generated 5′ overhang (5′-CTTG) was performed as described previously (50).

## ACKNOWLEDGEMENTS

The authors would like to thank members of the Landick lab for helpful discussions and Michael Engstrom and Michael Wolfe for comments on the manuscript. This work was supported by a grant the National Institutes of Health to R.L. (GM38660).

## Abbreviations

AED: 2-aminoethyl disulfide
cryo-EM: cryogenic electron microscopy
CTR: Cys-triplet reporter
CPR: Cys-pair reporter
DTT: dithiothreitol
EC: elongation complex
EDTA: ethylenediaminetetraacetic acid
IPTG: isopropyl β-D-1-thiogalactopyranoside
NAC: nucleotide addition cycle
NTP: nucleoside triphosphate
PAGE: polyacrylamide gel electrophoresis
PEC: paused elongation complex
ePEC: elemental paused elongation complex
hsPEC: hairpin-stabilized paused elongation complex
PEI: polyethylenimine
PMSF: phenylmethylsulfonyl fluoride
RH: rim helices
RNAP: RNA polymerase
SBHM: sandwich-barrel hybrid motif
SDS: sodium dodecyl sulfate
SI1: sequence insertion 1
SI3: sequence insertion 3
SPB: SI3 positional bias
TH: trigger helix
TL: trigger loop
TMAD: tetramethyldiazenedicarboxamide

## Supplementary Information for

This PDF file includes

1. Expanded Materials and Methods
2. SI Tables Table S1. Oligonucleotides and plasmids used in this study. Table S2. Kinetic fitting parameters for pause assays shown in Figs. 5 and 6. Table S3. Bacterial species used to screen for and align trigger-loop sequence insertions.
3. SI Figures Figure S1. Alternative locations of SI3 in ECs and PECs and CTR Cys residues. Figure S2. Optimization of the CTR crosslink assay using H_2_O_2_. Figure S3: H_2_O_2_ minimally affects pause kinetics of WT RNAP at the *his* elemental and hairpin-stabilized pause site. Figure S4. RNA extension assays for RNA–DNA scaffolds used in CTR assays in Figs. 2, 3, and 4. Figure S5. Ligated-scaffold transcription assay. Figure S6. Evolution of TL insertions in RNAP.
4. SI References

## Expanded Materials and Methods

### Materials

DNA and RNA oligonucleotides were purchased from Integrated DNA Technologies (IDT; Coralville, IA) and purified by 15% denaturing polyacrylamide gel electrophoresis (PAGE) before use. [*γ*-^32^P]ATP, [*α*-^32^P]CTP and [*α*-^32^P]GTP were obtained from PerkinElmer Life Sciences; rNTPs, from Promega (Madison, WI, USA); and 3′deoxy ATP (3′dATP) and CMPCPP, from Jena Bioscience. H_2_O_2_ 30% (w/w) solution and standard buffers and reagents were from Sigma-Aldrich or Thermo-Fisher.

### RNAP purification

RNAP expression plasmids were transformed into BL21(DE3) and a single colony was inoculated into 5 mL LB+50 µg kanamycin/mL and grown overnight growth at 37 °C. The saturated cell culture was then added to 1 L fresh LB+50 µg kanamycin/mL and grown at 37 °C with adequate aeration by orbital shaking in a Fernbach flask until apparent OD_600_ reached 0.6. Protein expression was induced by adding IPTG (Gold Biotechnology) to 1 mM and cell growth was continued for 3 hours. The cells were harvested and homogenized by sonication in 30 mL Lysis Buffer (50 mM Tris-HCl pH 7.9, 5% v/v glycerol, 233 mM NaCl, 2 mM EDTA, 10 mM β- mercaptoethanol, 10 mM DTT, 100 µg/mL PMSF, and 1 tablet of protease inhibitor cocktail (Roche cOmplete, Mini, EDTA-free). After removing cell debris by centrifugation (11000 × *g*, 15 min, 4 °C). DNA binding proteins including target RNAPs were precipitated by addition of polyethylenimine (PEI, Sigma-Aldrich) to 0.6% (w/v) final. After centrifugation (11000 × *g*, 15 min, 4 °C), the protein pellet was resuspended in 25 mL PEI Wash Buffer (10 mM Tris-HCl pH 7.9, 5% v/v glycerol, 0.1 mM EDTA, 5 µM ZnCl_2_, 500 mM NaCl) to remove non-target proteins. After centrifugation (11000 × *g*, 15 min, 4 °C), RNAP was eluted from the pellet into 25 ml PEI Elution Buffer (10 mM Tris-HCl pH 7.9, 5% v/v glycerol, 0.1 mM EDTA, 5 µM ZnCl_2_, 1 M NaCl). The crude extract of RNAPs was subjected to sequential FPLC purifications of Ni^2+^ column (HisTrap FF 5 ml, GE Healthcare Life Sciences) and heparin column (Heparin FF 5 ml, GE Healthcare Life Sciences) and the purified RNAP was dialyzed into RNAP Storage Buffer (10 mM Tris-HCl, 25% v/v glycerol, 100 mM NaCl, 100 µM EDTA, 1 mM MgCl_2_, 20 µM ZnCl_2_, 10 mM DTT) to final concentrations of 5–10 mg/mL. The high concentration of DTT (10 mM) was included to inhibit disulfide bond formation by engineered cysteine residues.

### CTR crosslinking assay, gel quantitation, and SPB calculation

The DNA–RNA bubble scaffold used for EC reconstitution was assembled by incubation purified oligos (20 µM non-template strand DNA, 15 µM template strand DNA, 10 µM RNA, 20 mM Tris-OAc pH 7.7, 5 mM Mg(OAc)_2_, 40 mM KOAc) in a thermal cycler (95 °C for 2 min, 75 °C for 2 min; 45-to-25 °C gradient over 25 min). To reconstitute CTR ECs, the CTR core RNAP was incubated with the scaffold at 1:2 ratio (2.5 µM CTR-RNAP, 5 µM scaffold) in 10 µL of Elongation Buffer (25 mM HEPES-KOH pH 8.0, 130 mM KCl, 5 mM MgCl_2_, 0.15 mM EDTA, 5% v/v glycerol, 25 µg acetylated BSA/ml) for 20 min at 37 °C. For reconstitution of EC with a complementary scaffold, only the RNA and template strand DNA were annealed (15 µM template strand DNA, 10 µM RNA) using the same solution and thermal conditions as for the bubble scaffold. The CTR core enzyme was incubated with the scaffold for 10 min at 37 °C, then non-template strand DNA was added to 20 µM and incubation was continued for 10 min. To mimic an RNA hairpin structure at the RNA exit channel, asRNA base pairing to nascent RNA was added to 25 µM and incubation was continued for 5 min.

For assays that involved nucleotide addition to reconstituted ECs prior to disulfide formation (Figs. 3, 4, and 5), NTPs were added to 50 µM each and the reactions were incubation at 37 °C for 5 min. Nucleotide incorporation was tested for completion by measuring the change in size of 5′ end ^32^P-labeled RNA by denaturing PAGE (Fig. S3).

For PAGE analysis, ∼0.3 uL samples were mixed with NuPAGE 4X LDS Sample Loading Buffer (Novex, reducing agents omitted) and electrophoresed through a thin layer 4-15% gradient polyacrylamide gel (GE Healthcare) using a PhastSystem Electrophoresis unit (originally from Pharmacia). Gels were stained with Imperial Protein Stain (Thermo Scientific), destained with water and imaged with a CCD camera (Protein Simple).

For comparison of crosslink efficiencies among oxidants, cystamine or diamide were used at 10 mM or 5 mM, respectively, and were quenched with 12.5 mM iodoacetamide. CuSO_4_ was used at 5 µM and was not quenched prior to PAGE.

We used the Fiji-a software package based on ImageJ to quantify the percentage of crosslinking. The intensity of two crosslinked bands (I_β-β′_ for SI3–SI1crosslink, I_β′-β′_ SI3–RH crosslink) and the uncrosslinked β and β′ bands (I_unxlink_) were measured. Because the SI3–SI1 crosslink was inter-subunit and the SI3–RH crosslink was intra-subunit, the two crosslink bands were easily distinguished (since MW_β_=151 kDa, MW_β′_=155 kDa, we assumed Coomassie brilliant blue G-250 binds β and β′ to similar levels). Thus, we calculated the % crosslinking from the band intensities (I) in the gel images as:

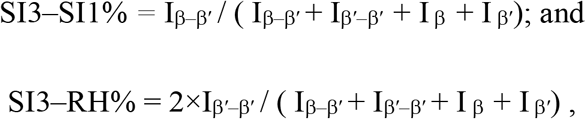

where the crosslinked species are indicated by subscripts containing two subunits and the intensity of the SI3–RH crosslinked species is doubled because it is approximately half the size of the SI3–SI1 crosslinked species. The SPB value was then calculated as the ratio between the two crosslink species:

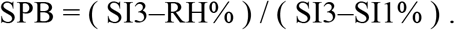

All SPB values were determined from at least 3 independent assays and reported as mean ± SD.

### Ligation-elongation assay and quantification

Pre-assembled scaffold (2 µL of 5 µM RNA, 10 µM template strand DNA) was incubated with 4 µL of CPR core enzyme (12.5 µM) in 20 µL Elongation Buffer (final RNA:RNAP ratio of 1:5; 0.5 µM RNA, 1.0 µM template strand DNA, 2.5 µM CPR RNAP) for 15 min at 37 °C. Nontemplate strand DNA (1.5 µL to 1.5 µM final concentration) was added and the 10 µL mixture was incubated for an additional 15 min at 37 °C. The estimated concentration of assembled EC was 500 nM.

To ligate the EC to the 2.2 kb DNA template, 270 nM EC was incubated with 90 nM purified DNA template, 1 × NEB T4 ligase buffer (including 1 mM ATP and 10 mM DTT) and 400 U T4 DNA ligase in a 10 µL reaction at 16 °C for 2 hours. The estimated concentration of ligated EC was 90 nM. To extend EC to the desired register and form the crosslinked EC, 50 nM ligated EC was radiolabeled by incubation with 2 µCi [*α*-^32^P] GTP (3000 Ci/mmol) for 2 min, extended with 100 µM each ATP and GTP for 10 min to form A26 EC, and then crosslinked with 5 mM H_2_O_2_ in a 20 µL reaction as described above. The mixture was then diluted with Elongation Buffer to 100 µL. The estimated final concentration of EC was 10 nM. To re-start elongation, 25 µL of 2 mM each NTP was combined with 25 µL ∼10 nM EC at 37 °C. Reaction samples (5 µL) were withdrawn at 10, 20, 30, 40, 60, 120 and 300 seconds and mixed with an equal volume of Stop Buffer (8 M urea, 50 mM EDTA, 90 mM Tris-borate buffer, pH 8.3, 0.02% bromophenol blue, 0.02% xylene cyanol). For rate measurements, triplicates of 20-s reactions were performed. RNAs in all collected samples were heated to 95 °C for 2 min before size separation by denaturing PAGE (8%, 19:1 acrylamide:bis-acrylamide, 45 mM Tris-borate, pH 8.3, 1.25 mM Na_2_EDTA, 8 M urea). The RNAs were run alongside a radiolabeled, MspI-digested pBR322 DNA ladder. The gel was exposed to a Storage Phosphor Screen (GE Healthcare) and scanned with Typhoon PhosphorImager.

The gels were quantified using ImageQuant software (GE Healthcare). To estimate the averaged elongation rates, the intensities of RNAs longer than 66 nt after 20 s extension with 1 mM NTPs (a 66 nt band was generated by unligated ECs) were determined and average elongation rates were calculated from the average sizes of these RNAs in 3 independent assays.

### *In vitro* transcription pause assays and quantification

For a typical pause assay, the annealed scaffold prepared as described above was incubated with CPR core enzyme with 1: 3 ratio (1 µM RNA, 2 µM template strand DNA, 3 µM CPR RNAP) in 60 µL Elongation Buffer for 15 min at 37 °C. For complementary scaffolds, non-template DNA was added to 5 µM and incubation continued for an additional 15 min. Further binding by CPR RNAP was blocked by addition of heparin to 0.1 mg/mL and 5 min incubation. To induce disulfide bond formation, 5 mM H_2_O_2_ was added and the mixture was incubated for 15 min at 37 °C. Samples with and without H_2_O_2_ addition were taken for PAGE analysis to measure crosslink efficiency in each experiment. Excess H_2_O_2_ was eliminated by addition of the mixture catalase to 0.1 U/mL followed by 5 min incubation at room temperature. The mixture was then diluted with 4 volumes of Elongation Buffer and radiolabeled by incubation with 2 µCi [*α*-^32^P]CTP (3000 Ci/mmol). To form paused ECs, CTP and UTP (2 µM each final) were added sequentially followed by 5 min incubation at 37 °C. The hairpin-stabilized paused complex was formed by addition of 1 µM asRNA at this point to reactions containing 200 nM ePEC and incubation was continued for 10 min. The PECs were then incubated at 37 °C for 2 min before initiating the pause assay by combination of equal volumes of 200 nM PEC solution and 20 µM GTP in Elongation buffer (100 nM ePEC/hsPEC and 10 µM GTP final). Samples (5 µL) were removed at 5, 10, 15, 30, 60 and 120 s and mixed with an equal volume of Stop Buffer. Before addition of GTP, a zero time point sample was generated by combination of 5 uL 200 nM PEC, 5 µL Elongation Buffer and 10 µL Stop Buffer. A final portion (4.45 µL) of PECs remaining at end of the time course was chased by addition of 0.55 µl 15 mM GTP and incubation for 1 min at 37 °C followed by addition of 5 µL Stop Buffer to assess the activity of the PECs. RNAs were analyzed by denaturing PAGE (15% 19:1 acrylamide: bis-acrylamide, 45 mM Tris-borate, pH 8.3, 1.25 mM Na_2_EDTA, 8 M urea). Gels were exposed to a Storage Phosphor Screen (GE Healthcare) and scanned with Typhoon PhosphorImager. Deviations from this protocol, where appropriate, are noted in the figure legends.

Pause assays were quantified using ImageQuant software (GE Healthcare). For all time point samples, the paused and the escaped RNA intensities were measured separately and the shorter transcripts corresponding to degraded RNA were not included in analysis. To correct for any readthrough transcripts formed before pause reaction started, these RNA intensities at time point zero were subtracted from each time point. In the similar way, the paused RNA intensity of the chased sample was subtracted from each time point to correct the inactive ECs/PECs. The corrected pause RNA as a function of time was used to determine pausing kinetics fitting to bi-exponential decay functions (1). (2)

### Protein sequence alignment

Seventy-two bacterial species (Table S2) were chosen from 38 representative phyla and the sequences of their β′ subunits were obtained from the National Center for Biotechnology Information protein database (NCBI, National Library of Medicine, Bethesda, MD; https://www.ncbi.nlm.nih.gov/). The β′ subunit sequences were checked for the existence of trigger loop sequence insertion. Thirty sequences containing trigger loop insertions were aligned with MUSCLE algorithm (3) and the alignment were transformed into sequence logos with WebLogo (4).

---

**Table S1.**
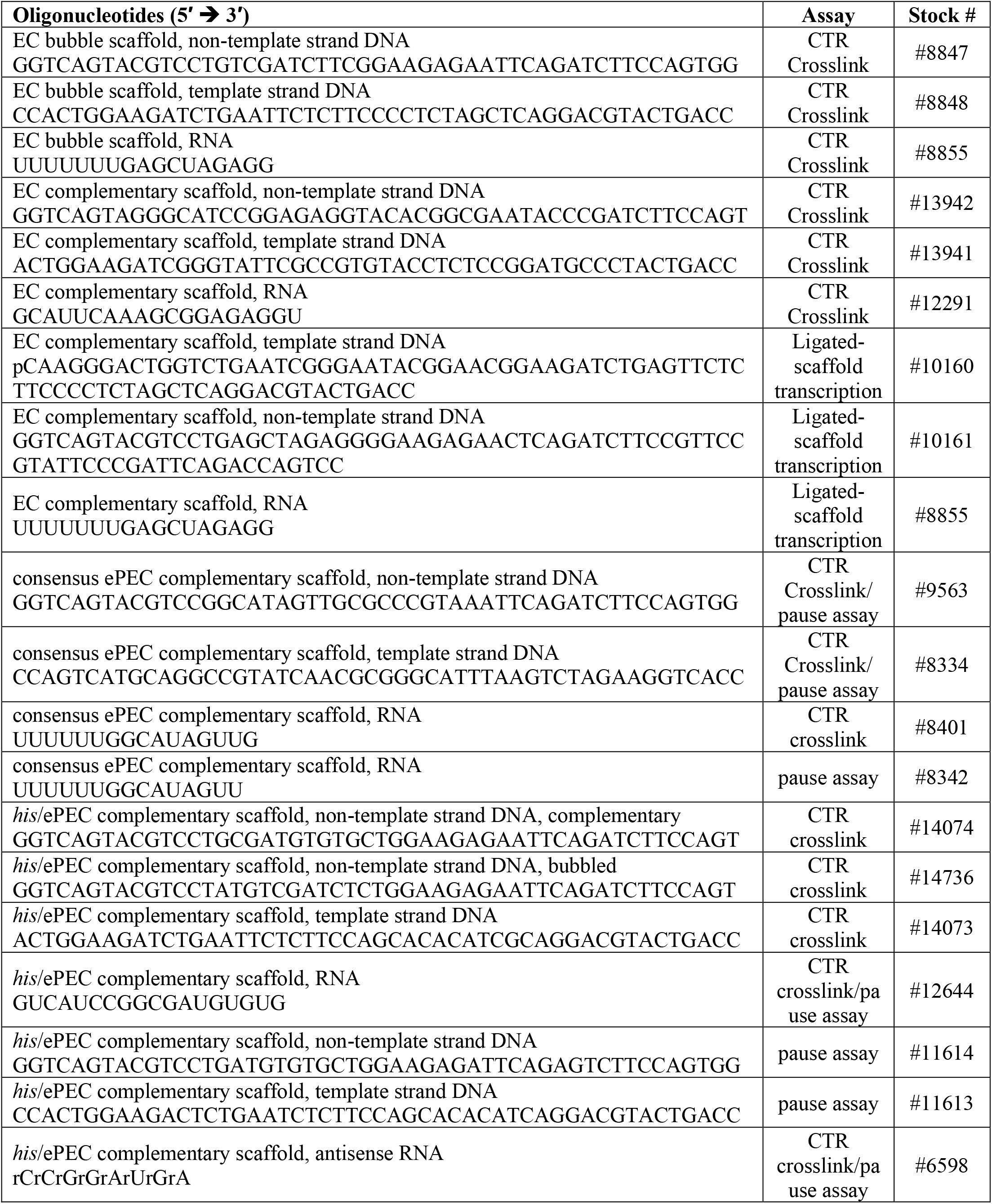
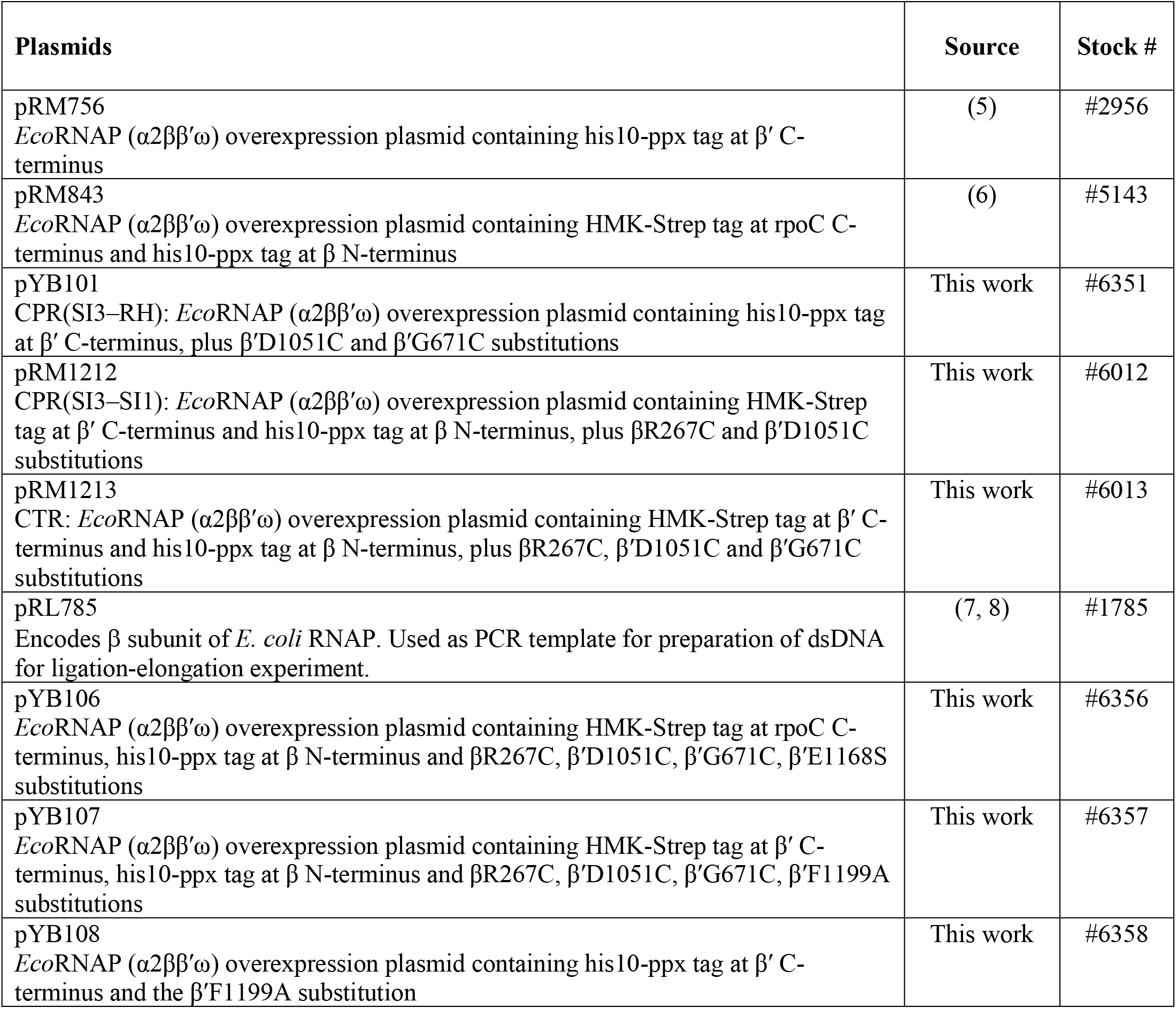
Oligonucleotides and plasmids.

**Table S2.** Kinetic fitting parameters for pause assays shown in Figs. 5 and 6. Fig. 5C.

| CPR(SI3-SI1) | $f_{\text{fast}}$ | error | $k_{\text{fast}}(\text{s}^{-1})$ | error | $f_{\text{slow}}$ | error | $k_{\text{slow}}(\text{s}^{-1})$ | error |
| --- | --- | --- | --- | --- | --- | --- | --- | --- |
| ePEC, red | 0.94 | 0.020 | 0.75 | 0.13 | 0.056 | 0.017 | 0.030 | 0.016 |
| ePEC, xlink | 0.34 | 0.0091 | 0.54 | 0.078 | 0.66 | 0.0059 | 0.0046 | 0.00018 |
| hsPEC, red | 0.28 | 0.028 | 0.28 | 0.060 | 0.72 | 0.026 | 0.022 | 0.0012 |
| hsPEC, xlink | 0.17 | 0.047 | 0.38 | 0.35 | 0.83 | 0.031 | 0.0031 | 0.00066 |

| CPR(SI3-RH) | $f_{\text{fast}}$ | error | $k_{\text{fast}}(\text{s}^{-1})$ | error | $f_{\text{slow}}$ | error | $k_{\text{slow}}(\text{s}^{-1})$ | error |
| --- | --- | --- | --- | --- | --- | --- | --- | --- |
| ePEC, red | 0.94 | 0.0086 | 0.74 | 0.064 | 0.058 | 0.0070 | 0.022 | 0.0052 |
| ePEC, xlink | 0.92 | 0.020 | 0.59 | 0.076 | 0.083 | 0.014 | 0.011 | 0.0048 |
| hsPEC, red | 0.44 | 0.092 | 0.22 | 0.080 | 0.56 | 0.090 | 0.024 | 0.0050 |
| hsPEC, xlink | 0.92 | 0.0077 | 0.59 | 0.031 | 0.083 | 0.0056 | 0.011 | 0.0020 |

| PECs | $f_{\text{fast}}$ | error | $k_{\text{fast}}(\text{s}^{-1})$ | error | $f_{\text{slow}}$ | error | $k_{\text{slow}}(\text{s}^{-1})$ | error |
| --- | --- | --- | --- | --- | --- | --- | --- | --- |
| ePEC, WT | 0.97 | 0.0090 | 0.73 | 0.070 | 0.030 | 0.0056 | 0.0047 | 0.0039 |
| ePEC, F1199A | 0.98 | 0.0029 | 0.86 | 0.042 | 0.018 | 0.016 | 0 | 0.0013 |
| hsPEC, WT | 0.55 | 0.12 | 0.13 | 0.031 | 0.44 | 0.13 | 0.026 | 0.0065 |
| hsPEC, F1199A | 0.76 | 0.066 | 0.31 | 0.043 | 0.24 | 0.065 | 0.044 | 0.013 |

| WT PEC | $f_{\text{fast}}$ | error | $k_{\text{fast}}(\text{s}^{-1})$ | error | $f_{\text{slow}}$ | error | $k_{\text{slow}}(\text{s}^{-1})$ | error |
| --- | --- | --- | --- | --- | --- | --- | --- | --- |
| ePEC, red | 0.91 | 0.079 | 1.0 | 1.7 | 0.089 | 0.077 | 0.067 | 0.064 |
| ePEC, xlink | 0.96 | 0.071 | 1.3 | 6.4 | 0.040 | 0.070 | 0.067 | 0.13 |
| hsPEC, red | 0.22 | 0.017 | 0.43 | 0.11 | 0.78 | 0.014 | 0.021 | 0.00073 |
| hsPEC, xlink | 0.34 | 0.029 | 0.50 | 0.16 | 0.66 | 0.026 | 0.029 | 0.0019 |

**Table S3.** Bacterial species used to screen for and align trigger-loop sequence insertions.

| Phylum | Genus Species | Accession | $\beta'$, aa | SI3? | SI3, aa |
| --- | --- | --- | --- | --- | --- |
| Thermus | *Thermus thermophilus* | Q8RQE8 | 1524 | No | 0 |
|  | *Deinococcus geothermalis* | Q1J0P7 | 1537 | No | 0 |
|  | *Deinococcus radiodurans* | Q9RVW0 | 1546 | No | 0 |
| Thermotogae | *Petrotoga mobilis* | A9BF32 | 1643 | No | 0 |
|  | *Thermotoga maritima* | P36252 | 1690 | No | 0 |
|  | *Thermosipho melanesiensis* | A6LKB7 | 1649 | No | 0 |
| Aquificae | *Aquifex aeolicus* | O67763 | 1547 | Yes | 206 |
|  | *Balnearium lithotrophicum* | WP_142934069.1 | 1471 | Yes | 168 |
|  | *Hydrogenothermus marinus* | WP_121922216.1 | 1602 | Yes | 207 |
| Actinobacteria | *Mycobacterium tuberculosis* | P9WGY7 | 1316 | No | 0 |
|  | *Streptomyces avermitilis* | Q82DQ4 | 1299 | No | 0 |
|  | *Bifidobacterium adolescentis* | A1A316 | 1337 | No | 0 |
| Chloroflexi | *Dehalococcoides mccartyi* | Q3Z8V3 | 1295 | No | 0 |
|  | *Herpetosiphon aurantiacus* | A9B6J1 | 1543 | No | 0 |
|  | *Chloroflexus aurantiacus* | A9WH11 | 1503 | No | 0 |
| Cyanobacteria | *Gloeobacter violaceus* | Q7NDF7 | 1262 | Yes | 633 |
|  | *Synechococcus elongatus* | Q5MZ21 | 1318 | Yes | 638 |
|  | *Trichodesmium erythraeum* | Q110H3 | 1415 | Yes | 684 |
| Firmicutes | *Clostridioides difficile* | A0A125V165 | 1161 | No | 0 |
|  | *Bacillus subtilis* | P37871 | 1199 | No | 0 |
|  | *Mycoplasma pneumoniae* | A0A0H3DMP7 | 1290 | No | 0 |
| Tenericutes | *Acholeplasma equifetale* | WP_026399639.1 | 1367 | No | 0 |
|  | *Entomoplasma ellychniae* | PPE04721.1 | 1252 | No | 0 |
|  | *Haloplasma contractile* | ERJ12394.1 | 1204 | No | 0 |
| Planctomycetes | *Rhodopirellula baltica* | Q7URW4 | 1429 | Yes | 173 |
|  | *Chlamydia trachomatis* | Q3KM48 | 1396 | Yes | 190 |
|  | *Chlamydia abortus* | Q5L5I4 | 1393 | Yes | 190 |
| Spirochaetes | *Leptospira interrogans* | Q8F0S3 | 1404 | Yes | 223 |
|  | *Borrelia garinii* | B7XT60 | 1377 | Yes | 224 |
|  | *Treponema pedis* | S5ZS96 | 1424 | Yes | 228 |
| Bacteroidetes | *Chlorobium phaeobacteroides* | A1BD24 | 1492 | Yes | 190 |
|  | *Bacteroides fragilis* | Q5L898 | 1427 | Yes | 180 |
|  | *Flavobacterium johnsoniae* | A5FIJ4 | 1436 | Yes | 180 |
| δ-proteobacteria | *Sorangium cellulosum* | A9GRB4 | 1427 | Yes | 185 |
|  | *Geobacter sulfurreducens* | I7EPD8 | 1395 | Yes | 182 |
|  | *Desulfovibrio vulgaris* | Q727C6 | 1385 | Yes | 182 |
| ε-proteobacteria | *Wolinella succinogenes* | Q7MA56 | 2883 | Yes | 332 |
|  | *Sulfurovum lithotrophicum* | WP_046551950.1 | 1509 | Yes | 333 |
|  | *Campylobacter hominis* | WP_041570540.1 | 1509 | Yes | 331 |
| α-proteobacteria | *Ehrlichia ruminantium* | Q5HC04 | 1411 | Yes | 186 |
|  | *Zymomonas mobilis* | A0A0H3FXE3 | 1391 | Yes | 178 |
|  | *Caulobacter vibrioides* | Q9AAU1 | 1396 | Yes | 183 |
| β-proteobacteria | *Acidovorax delafieldii* | C5T742 | 1408 | Yes | 188 |
|  | *Herminiimonas arsenicoxydans* | A4G9U4 | 1414 | Yes | 189 |
|  | *Bordetella petrii* | A9IJ22 | 1412 | Yes | 188 |
| γ-proteobacteria | *Legionella pneumophila* | Q5ZYP9 | 1401 | Yes | 188 |
|  | *Aliivibrio fischeri* | Q5E239 | 1401 | Yes | 188 |
|  | *Escherichia coli* | P0A8T7 | 1407 | Yes | 188 |
| Zetaproteo | *Mariprofundus ferrooxydans* | WP_018295044.1 | 1399 | Yes | 189 |
| Oligoflexia | *Bdellovibrio bacteriovorus* | WP_063244102.1 | 1376 | Yes | 183 |
| Chrysiogenetes | *Chrysiogenes arsenatis* | WP_084417671.1 | 1347 | Yes | 175 |
| Acidobacteriia | *Acidobacterium capsulatum* | C1F3Y4 | 1389 | Yes | 173 |
| Blastocatellia | *Pyrinomonas methylaliphatogenes* | WP_041973492.1 | 1389 | Yes | 173 |
| Dictyoglomia | *Dictyoglomus thermophilum* | WP_012547665 | 1429 | Yes | 258 |
| Synergistia | *Aminomonas paucivorans* | WP_006301288.1 | 1654 | Yes | 314 |
| Thermodesulfobacteria | *Thermodesulfobacterium commune* | WP_038061948.1 | 1353 | Yes | 183 |
| Nitrospira | *Nitrospira defluvii* | CBK41054.1 | 1396 | Yes | 197 |
| Deferribacteres | *Caldithrix abyssi* | APF17647.1 | 1429 | Yes | 175 |
| Fusobacteriia | *Fusobacterium necrophorum* | RXZ69643.1 | 1322 | Yes | 165 |
| Gemmatimonadetes | *Gemmatimonas aurantiaca* | BAH37898.1 | 1436 | Yes | 182 |
| Fibrobacteres | *Fibrobacter succinogenes* | WP_173384178.1 | 1480 | Yes | 178 |
| Verrucomicrobia | *Verrucomicrobium spinosum* | WP_009964681.1 | 1376 | Yes | 174 |
| Lentisphaerae | *Lentisphaera araneosa* | WP_007279810.1 | 1423 | Yes | 174 |
| Kiritimatiellaeota | *Kiritimatiella glycovorans* | AKJ64962.1 | 1383 | Yes | 174 |
| Chlamydiae | *Chlamydia abortus* | ASD30784.1 | 1393 | Yes | 190 |
| Endomicrobia | *Endomicrobium proavitum* | WP_052571414.1 | 1580 | Yes | 184 |
| Elusimicrobia | *Elusimicrobium minutum* | WP_012415591 | 1385 | Yes | 173 |
| Armatimonadetes | *Armatimonas rosea* | WP_184203436.1 | 1473 | Yes | 52 |
| Acidithiobacillia | *Acidithiobacillus caldus* | AUW32002.1 | 1382 | Yes | 189 |
| Candidate phyla radiation | Parcubacteria | KND51703.1 | 1253 | No | 0 |
| Candidate phyla radiation | Microgenomates | RJR29364.1 | 1257 | No | 0 |

**Fig. S1.**
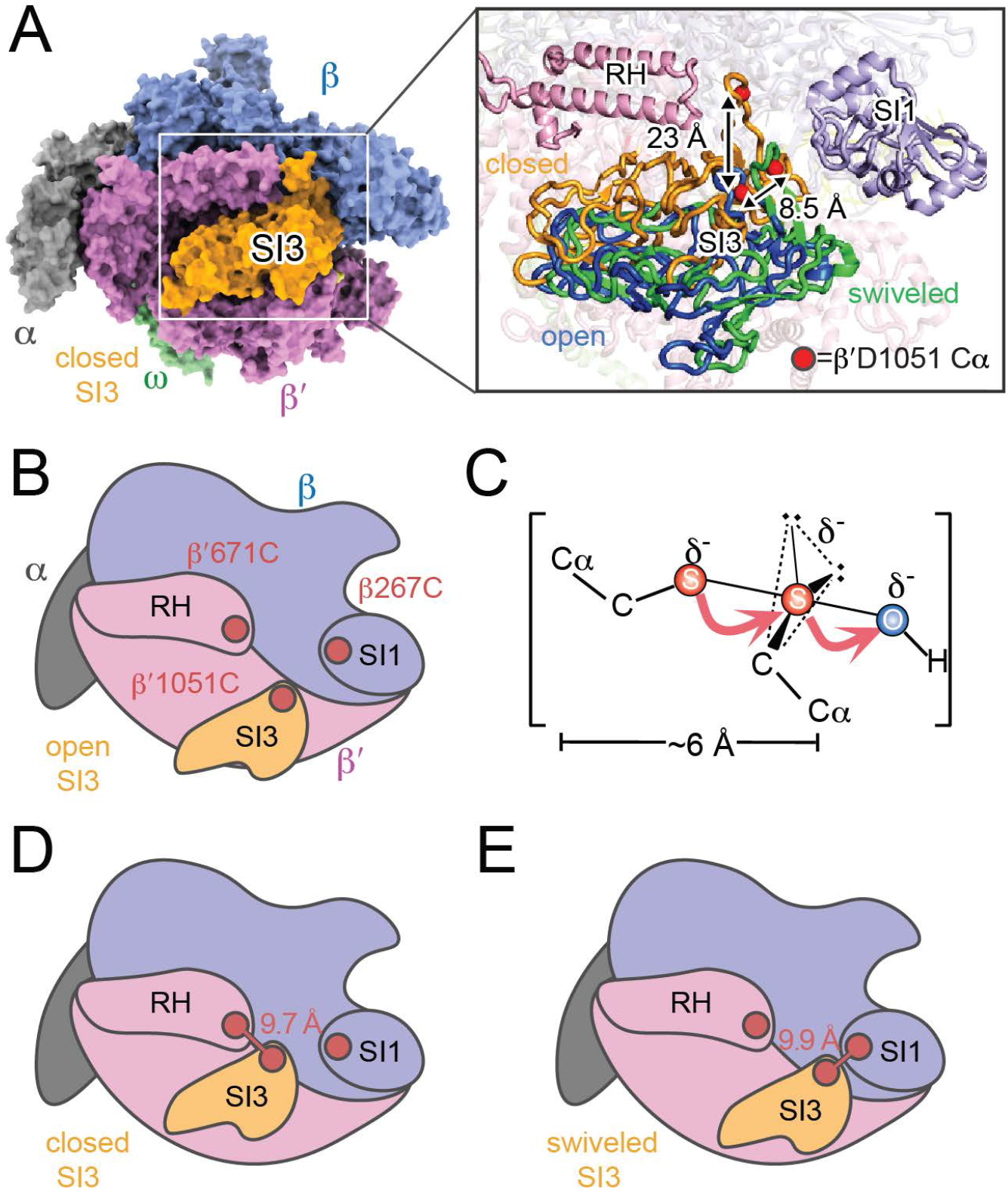
Alternative locations of SI3 in ECs and PECs and CTR Cys residues. (*A*) Locations of SI3 in ECs, RNAP is shown in the TH conformation with SI3 in closed position (pdb 4yln) (9). Inset, superposition of three different SI3 locations: closed (orange, pdb 4yln); open (blue, pdb 6alf), and swiveled (green, pdb 6asx) (2, 9, 10). (*B*) Schematic representation of CTR. (*C*) Reaction geometry of disulfide formation with H_2_O_2_. (*D*), (*E*) C*α*–C*α* distances of residues changed to Cys to generate the CTR (red spheres): β′SI3–β′RH (closed) and β′SI3–βSI1 (swiveled).

**Fig. S2.**
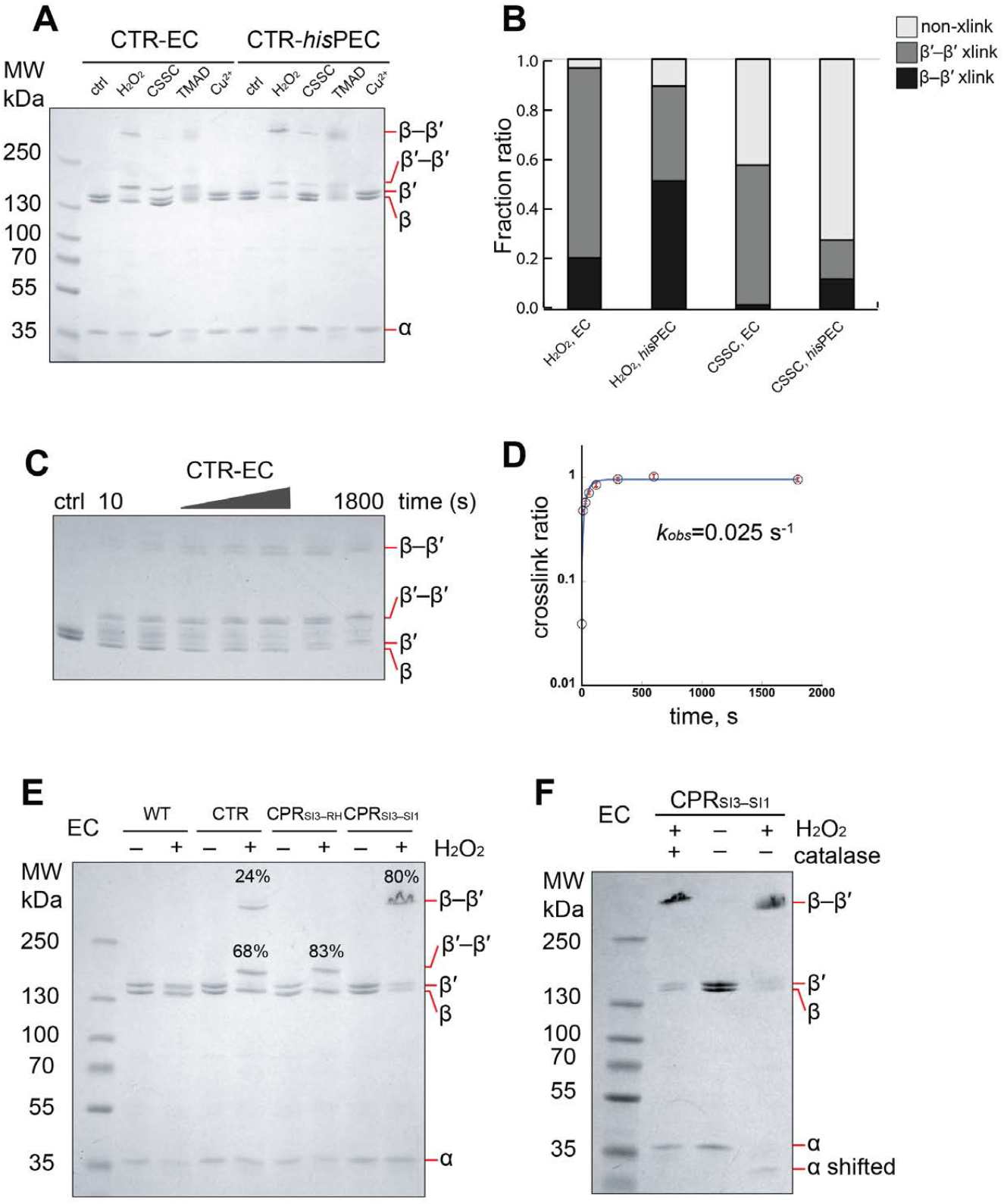
Optimization of the CTR crosslink assay using H_2_O_2_. (A) CTR crosslink assay of in EC and *his*PEC with different oxidants. CSSC, cystamine; TMAD, diamide. See scaffold sequences in Figs. 3 and 4. (B) Fractions of crosslinked and uncrosslinked species produced by cystamine or H_2_O_2_. (C) Time course of H_2_O_2_-mediated crosslinking in CTR RNAP reconstituted on an EC scaffold. (D) Plot of crosslink ratio as a function of time for CTR RNAP. Fit to a single exponential gives a pseudo-first-order rate of 0.025 s^−1^. (E) Comparison of crosslink species formed by CTR, CPR_SI3–RH_, and CPR_SI3-SI1_ RNAPs. (F) Removal of excess H_2_O_2_ by catalase treatment does not affect crosslink ratio but eliminates a shift in *α* subunit electrophoretic mobility.

**Fig. S3.**
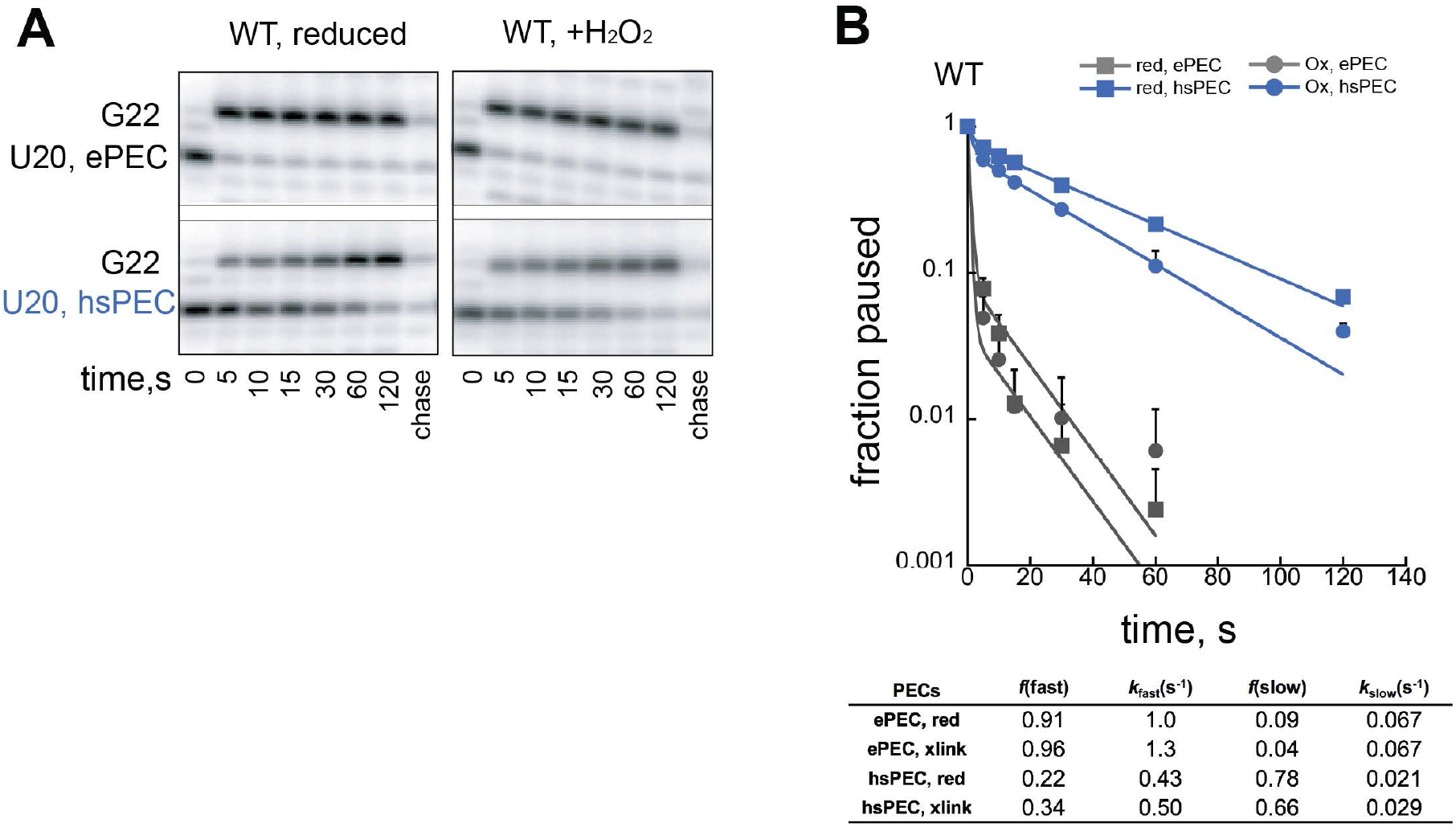
H_2_O_2_ minimally affects pause kinetics of WT RNAP at the *his* elemental and hairpin-stabilized pause site. (*A*) Pause escape by *his* ePEC and hsPEC in reducing and oxidizing conditions. (*B*) Pause kinetics of *his* ePEC and hsPEC in reducing and oxidizing conditions.

**Fig. S4.**
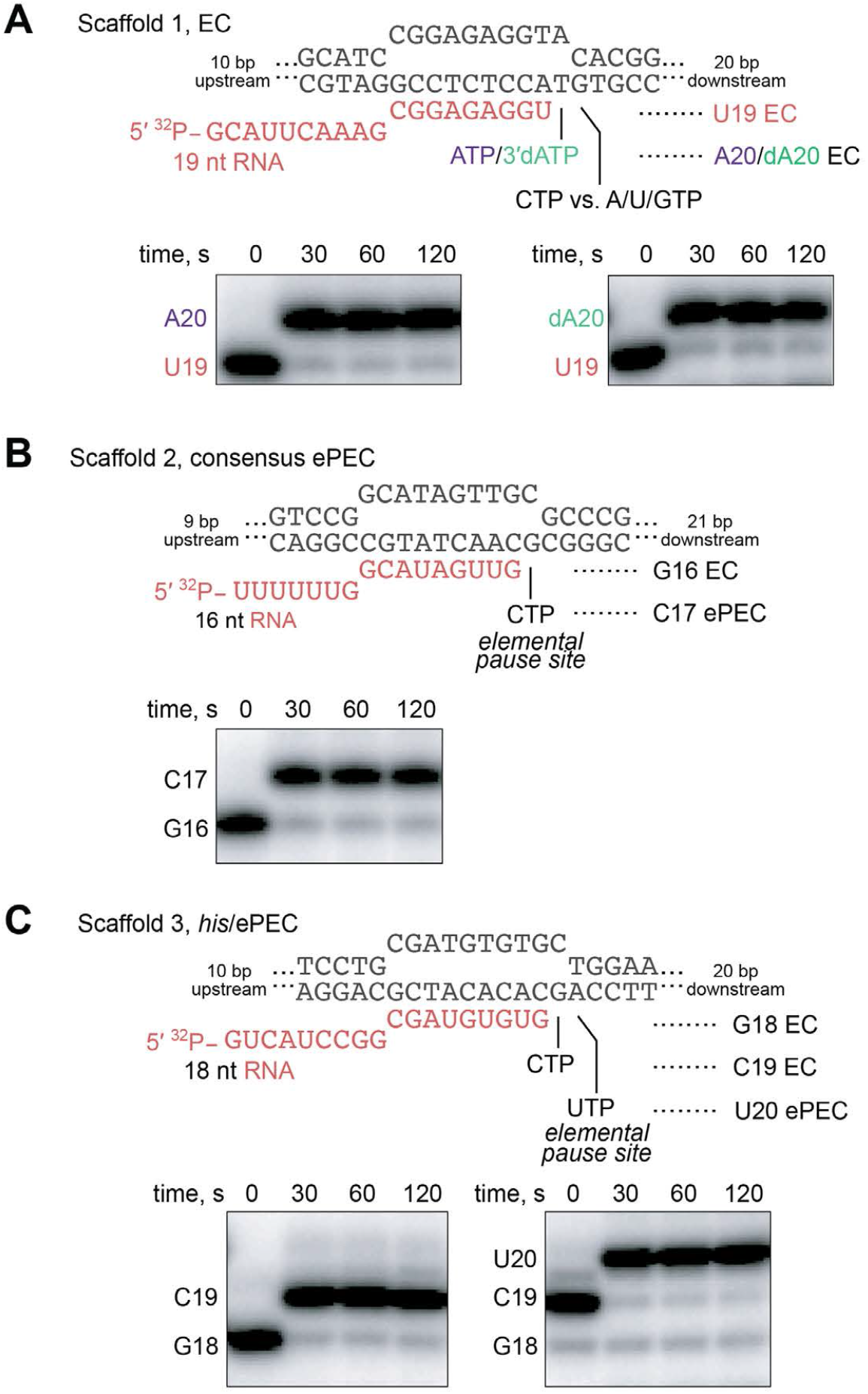
RNA extension assays for RNA–DNA scaffolds used in CTR assays in Figs. 2, 3, and 4. The 5′ ends of the RNAs were radiolabeled prior to reconstitution ECs and then reacted with 100 µM cognate NTPs. Samples were analyzed by denaturing PAGE (Materials and Methods) to verify that the desired nucleotide incorporation was complete in the CTR assays.

**Fig. S5.**
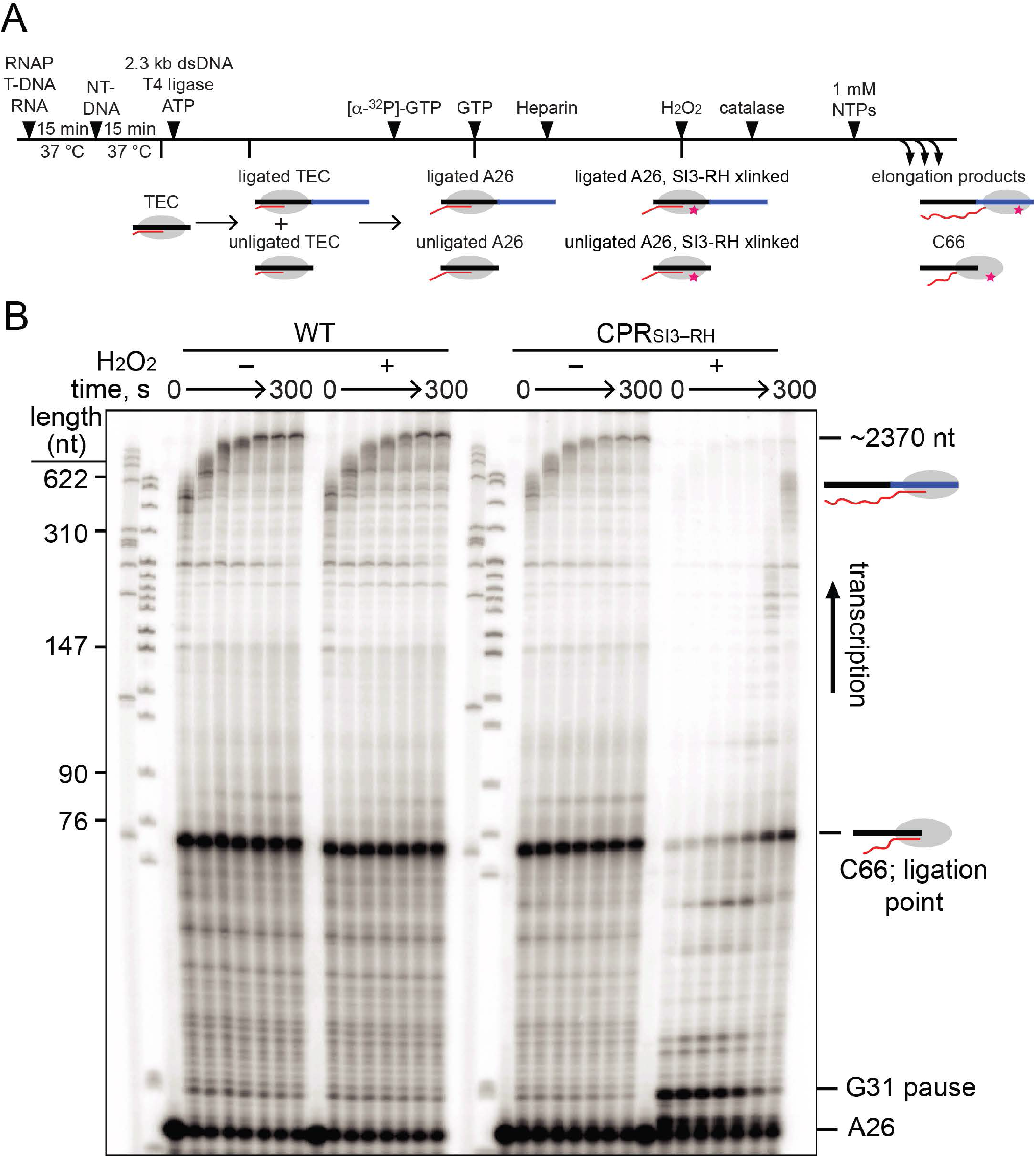
Ligated-scaffold transcription assay. (*A*) Schematic of ligated-scaffold transcription assay. (*B*) Phosphorimage of denaturing gel separating RNAs formed during the ligated-scaffold transcription assay by WT RNAP and CPR_SI3–RH_ in either reducing and oxidizing conditions (*Materials and Methods*)

**Fig. S6.**
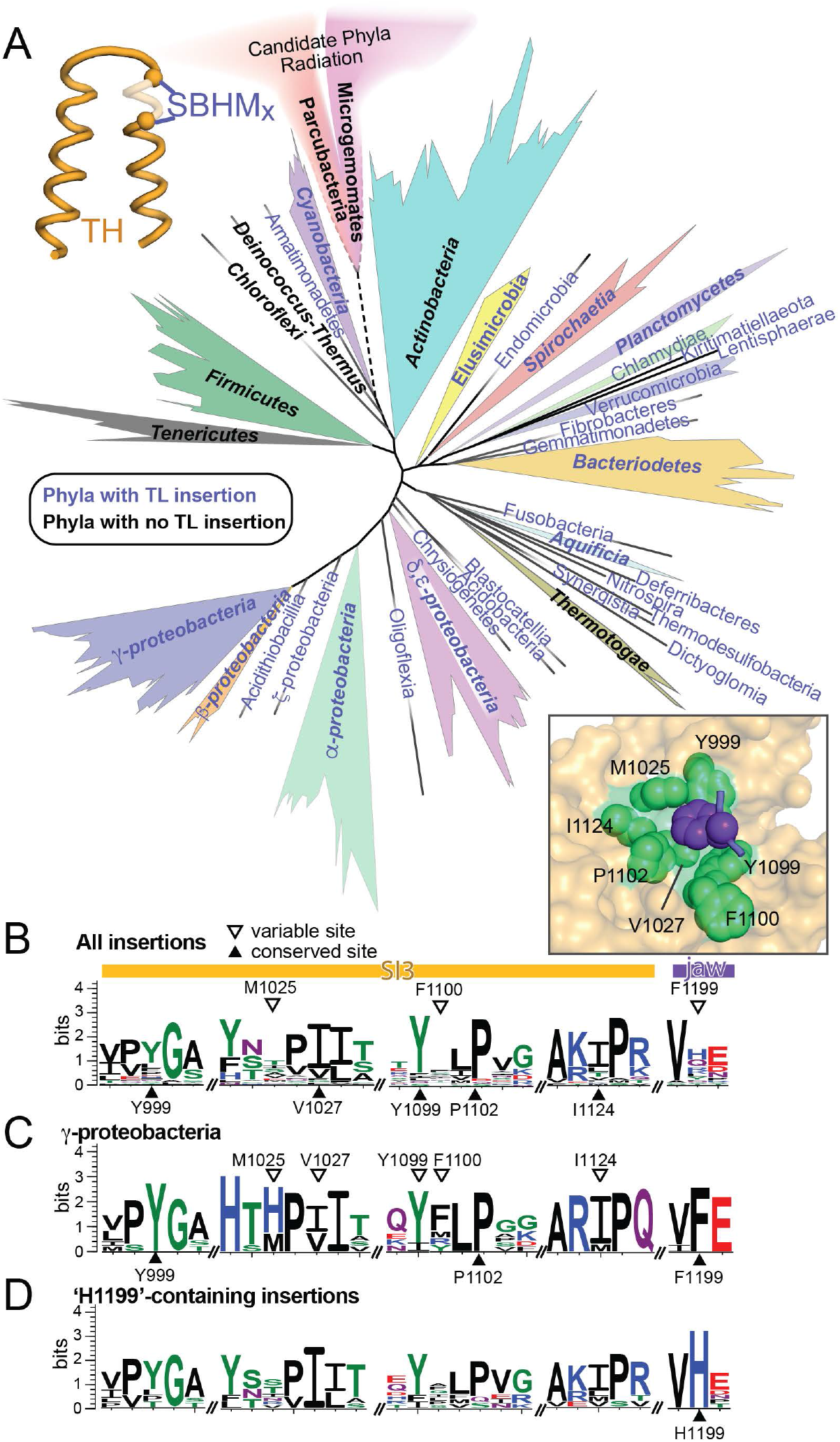
Evolution of TL insertions in RNAP. (*A*) An evolutionary tree depicting bacterial phylogeny based on 49 universally present genes (11) with lineages in which the TL contains an insertion of two or more SBHM domains indicated in blue. The area covered by each lineage is an approximate reflection of the number of known species in it. Conservation of residues involved in the Phe-pocket interaction: (*B*) in all TL insertion-containing bacteria; (*C*) in *γ*- proteobacteria; or (*D*) in TL insertion-containing bacteria in which the jaw Phe residue is replaced by His.

